# DipSkmer: Reference-free population genomics with diploid genome skims

**DOI:** 10.64898/2026.06.05.730460

**Authors:** Eduardo Charvel, Homère J. Alves Monteiro, Siavash Mirarab, Vineet Bafna

## Abstract

Ecologists and conservation biologists rely on genetic diversity as a key essential biodiversity variable (EBV) used to track population health and dynamics, and utilize the population parameter *θ* (estimated by the average pairwise genomic distance) as a key metric of diversity. While whole-genome-sequencing (wgs) is increasingly affordable, it will be considerable time before the full diversity of life is represented by high-quality assembled genomes; even then, constant monitoring will still require repeated sampling of populations. In contrast, genome skimming (low-coverage, short-read wgs) is highly cost-effective but challenging to analyze because the coverage is too low for assembly and reliable error correction. Mature methods, such as Mash, exist for estimating pairwise genomic distances based on the Jaccard similarity of *k*-mer sets computed using sketching techniques. Some, such as Skmer, additionally model the impacts of low coverage. These methods have been successfully applied to assembly-free species identification and phylogenetics; however, their use in population genetics has been limited. This is because these methods implicitly treat genomes as haploid and heterozygosity confounds true estimates of genomic distance for diploid organisms. In this paper, we address this problem through a number of technical advances. First, we use coalescent theory to mathematically derive how the Jaccard index between two diploid samples changes with the scaled population size parameter (*θ*). Next, we derive an estimator that computes *θ* from the Jaccard index, in addition to several auxiliary variables, which we also estimate from the genome skims. The resulting method, DipSkmer, enables more accurate estimates of coverage, sequencing error, and pairwise nucleotide distance for diploid samples. Analyses of both simulated and empirical datasets show that for diploids and low distances (e.g., *<* 2%), DipSkmer produces the most accurate pairwise distance estimates, outperforming existing alignment-free methods such as Mash and Skmer, and closely approximates ANGSD, a reference and alignment-based tool.

**Availability:** The code for DipSkmer is available at https://github.com/echarvel3/ReSkmer/tree/DipSkmer-REFACTOR. Simulation scripts and environments are available at https://github.com/echarvel3/dipskmer_scripts.

**Author Summary:** The process of obtaining full-genome population genomic measurements for biodiversity monitoring remains expensive due to the need for high-coverage sequencing and reference assemblies. Genome skimming has been shown to be a viable, low-coverage alternative for obtaining genomic distances, and alignment- and assembly-free methods exist for analyzing nuclear data from skimming data to estimate the distance between samples. However, existing methods fail to model within-sample heterozygosity, expected for diploid organisms. Given the dominance of diploidy among species of interest to ecologists, the implications of these simplifying assumptions warrant further study. Here, we present a mathematical model of the *k*-mer sets sampled from two diploid genomes from a Wright-Fisher population. We use the model to develop DipSkmer, a *k*-mer-based, reference-free method for estimating nucleotide diversity and population divergence that, unlike its predecessors, models within-sample heterozygosity. Benchmarking shows more accurate genomic diversity estimates compared to existing reference-free, genome-skimming methods and comparable performance to the popular high-coverage, reference-based method, ANGSD. Thus, DipSkmer enables accessible, less expensive population monitoring through genetic diversity estimates.

## 1 Introduction

Nucleotide diversity, measured as the percentage of positions in the genome that differ between two organisms, is widely used across biology. This distance measure is used in phylogenetics, microbiome analyses, comparative genomics, and population genetics as a simple measure of evolutionary divergence. In particular, nucleotide diversity is among the Essential Biodiversity Variables (EBV) used extensively by ecologists and conservation biologists to monitor the health of species^35^. Species with dwindling populations suffer from reduced genetic diversity^16^, which can lead to loss of fitness^41^. Monitoring the health of species, which is increasingly important as biodiversity continues to face unprecedented levels of strain^8,17^, requires not just a one-time estimate of diversity but continuous monitoring across spatial and temporal gradients. Such a goal can only be achieved if we have access to low-cost tools for monitoring.

When reference genomes and highly covered samples are available, nucleotide diversity can be measured by calling genotypes and computing the Hamming distance. Under the Wright-Fisher (WF) model, Hamming distance corresponds^29^ to *θ* = 2*µN*_*e*_ for haploid and *θ* = 4*µN*_*e*_ for diploid populations, where *µ* is the substitution rate per nucleotide per site, and *N*_*e*_, the effective population size, is what interests the ecologists. However, much of the biodiversity on earth is not yet represented with reference genomes, and conservation biologists cannot afford to wait until such sampling is done. Even when reference genomes are available, high-coverage samples are expensive to obtain if repeated sampling is required. The viable alternative of barcoding a single marker gene gives limited resolution at the species level^25,37^.

An alternative that has been put forward^4^ and used with some success^2,7,9,10,13,24,28,52^ is genome skimming, which consists of sequencing a sample at low coverage without assembling the nuclear genome (mitogenome can be assembled, a method many skimming-based analyses use^12^). Skimming can have a low cost and can eliminate the need for assembly. The *k*-mer sketching methods can then be used to efficiently compute distances between sets of reads for species identification. In particular, Ondov et al.^34^ developed an efficient approach to use min-hash^5^ to estimate the Jaccard (*J*) index between the *k*-mer sets of two samples, and used a simple transformation noted earlier by Fan et al.^14^ to translate the Jaccard index to nucleotide distances. Later, Sarmashghi et al.^43^ showed how the translation between *J* and distance can also account for effects of low coverage and sequencing error by pairing it with an analysis of the *k*-mer frequency spectrum, and Charvel et al.^11^ showed how to account for the impact of repeats. Many researchers have built on various hashing techniques for fast distance calculation^1,18,22,36,40,45^.

Despite this rich literature, existing methods have largely overlooked the complexity introduced by within-sample heterozygosity. These methods fail to account for changes between the two alleles present in a single sample of a diploid organism, and instead model evolutionary changes on one chromosome, effectively assuming haploidy. Such a model is suitable for application to microbial species, but insufficient for diploid organisms. Ignoring within-sample heterozygosity may be inconsequential when comparing samples from different species (e.g., for species identification), but it is important for population genetic analyses at low distances. The fundamental unanswered mathematical questions are: *Given reads generated from two diploid individuals sampled from a Wright Fisher evolving population with scaled population size θ* = 4*µN*_*e*_, *how does the Jaccard index between their k-mer sets relate to θ, and how does Jaccard change as coverage drops and sequencing errors are added?* We resolve these questions here.

First, we use coalescence theory to characterize the Jaccard index with respect to *θ* for diploid genomes (Methods 4.2). Next, we show how Jaccard changes given low-coverage genome skims (Methods 4.3), and how to estimate parameters needed for Jaccard computation (Methods 4.4). We use these derivations to develop DipSkmer (Diploid Skmer), which takes as input two sets of reads from closely related diploid organisms and outputs their nucleotide distance. In extensive simulations, DipSkmer shows substantially higher accuracy compared to Mash and Skmer for diploid samples. On real data, we show that DipSkmer’s *θ* estimates correlate strongly with those from high-coverage data mapped to a reference genome.

## 2 Results

### 2.1 Diploidy Alters the Jaccard Index’s Relationship with Nucleotide Diversity

When comparing *k*-mers between two diploid genome skimming samples, each sample carries its own set of heterozygous and homozygous *k*-mers, which are in turn homozygous or heterozygous across both samples, thereby creating a complex relationship between within-sample heterozygosity and *k*-mer set similarity (Fig. 1A). At the individual *k*-mer level, it is hard to know whether it is heterozygous in each sample. However, by assuming the Wright-Fisher model, we can model the expected number of *k*-mers following particular evolutionary patterns, and thereby characterize how genetic diversity shapes the Jaccard Index. In DipSkmer (Diploid Skmer), we use a coalescent model to capture how *k*-mers, treated as independent haplotype blocks, can have distinct evolutionary histories (Fig. 1B-C). Given the resulting *k*-mer alleles, we then model how they are represented in the Jaccard Index (Fig. 1D), and finally incorporate a coverage and error model to account for missing and erroneous *k*-mers within the diploid framework (Fig. 1E), yielding a final estimate of *θ* (Fig. 1F).

**Figure 1:**
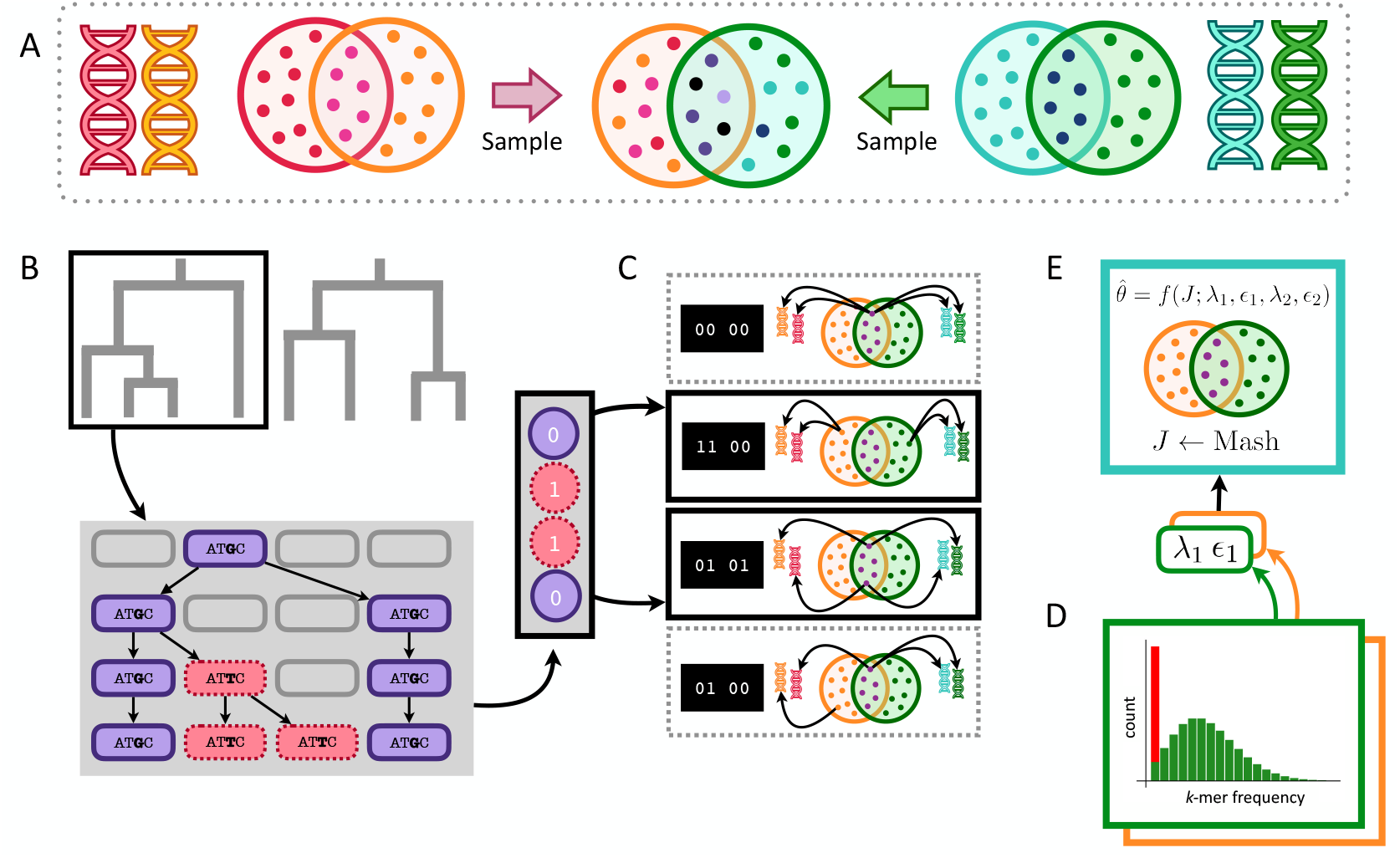
DipSkmer estimates nucleotide diversity between two diploid genomes, using only low coverage, error-prone, unassembled genomic data. **A)** The Jaccard index computed from genomes of two diploid organisms measures similarity between two pairs of haploid chromosomes, where each sample contains its own homozygous and heterozygous *k*-mers. **B)** We can model the possible coalescence scenarios of the four independent *k*-mers obtained from each position of two diploid genomes. **C)** For each topology, depending on the branches where mutations are introduced, *k*-mers may appear in one or both samples with computable probabilities (see Fig. 6). **D)** The alleles resulting from these mutations are then independently sorted into each haploid, leading to 0, 1, or 2 shared *k*-mers per position. **E)** We incorporate a model to estimate coverage (*ε*) and error (*θ*) based on *k*-mer frequency spectrum of the sample; erroneous *k*-mers (red) tend to be less frequent, whereas error-free *k*-mers tend to follow a Poisson distribution (green). **F)** We derive the expected numbers of *k*-mers shared in scenarios shown in (D) to design an estimator of nucleotide diversity (Eqs. (2) and (3)) given an observed Jaccard index and estimates of coverage and error.

#### Estimating *θ* between diploid samples

Under a diploid model, *k*-mer loci will occur twice in a DNA sample, making it so that each locus can weight the Jaccard differently with different configurations of heterozygous *k*-mers (Fig. 1D). DipSkmer’s nucleotide diversity model (Methods: Extending to two diploid individuals (*n* = 4)) assumes that all haplotypes are sampled from a randomly mating population, such that coalescence between the paternal and maternal alleles within a sample is identically distributed to coalescence between alleles from different individuals. Each set of haplotypes can then be traced back to one of two topologies, each with its own probability of occurrence (Fig. 1B). Under the infinite sites assumption, we model how mutations may fall on each topology (Fig. 1C) and assign a random permutation of haplotypes to the topology leaves (Fig. 1D). We then analyze the expected number of *k*-mers in the intersection (*I*) and union (*U*) at a single genomic position. Since different substitution patterns contribute differently to the union and intersection, enumerating all scenarios yields: 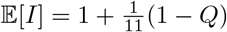 and 𝔼[*U*] = 2 − *Q*. Using the method of ratios from Mash/Skmer and solving for *Q*, we obtain:

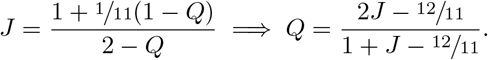

#### Diploid parameter estimation

To the diploid Jaccard model, we introduce a low-coverage model (Methods: Modeling low coverage and errors) that estimates the probability of a locus contributing an observed *k*-mer to the union and intersection of the Jaccard Index, given estimates of *k*-mer coverage (*λ*) and sequencing error rate (*ϵ*) (Fig. 1E). To estimate these sequencing parameters, we derive a new set of equations that account for how within-sample heterozygous *k*-mers affect the distribution of observed *k*-mer counts (Methods: Parameter estimation for diploid skims). Specifically, we take the ratio of consecutive *k*-mer count bins to obtain an error-free *k*-mer coverage estimate (*ξ*), which yields an internal, per-sample estimate of *θ* and, in turn, a *θ*-informed estimate of *λ* and sequencing error rate *ϵ*.

### 2.2 Simulation Results

In order to test DipSkmer’s accuracy, we use msprime^3^ to simulate diploid populations of different base genomes as as explicitly defined in Experimental setup. By modulating effective population size, simulated samples result in having different nucleotide diversity values (*θ* ∈ {0.001, 0.003, 0.006, 0.009, 0.012}). To test whether DipSkmer can also estimate the divergence between populations, we perform separate msprime simulations that lead to populations having different levels of genetic divergence (0.03125, 0.0625, 0.125, 0.25, 1) in coalescent units (*τ/*4*N*_*e*_). Note that these simulations violate the random mating assumptions of DipSkmer, and thus, test the robustness of the method (especially at higher divergence levels). Moreover, the presence of repeated *k*-mers violates DipSkmer assumptions; we focus on three base genomes with *k*-mer uniqueness above 0.7 in the main results. To test the degradation of performance with extremely high levels of repeats, we separately examine three extra genomes (Experimental setup). After generating genomes, we use ART^15^ to simulate reads for each simulation at varying coverages (*c* ∈ {0.5×, 1×, 2×, 4×, 8×}).

We compare DipSkmer to its predecessor alignment-free methods, Skmer (which effectively assumes haploid input) and Mash (which additionally ignores low coverage). We also compare to ReSkmer, which like Skmer ignores heterozygosity but unlike DipSkmer models repeats in the genome. We compare *θ* estimates using these methods to the measure of true *θ* (the percentage of substituted positions in genome). We measure error as 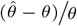, averaged over all replicates, where 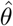 is the estimated pairwise distance and *θ* is the true pairwise distance. We also report mean absolute error (MAE), defined as 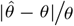. For population differentiation experiments, we estimate *D*_*xy*_ (average *θ* between pairs of samples taken from different populations) and Weir and Cockerham’s 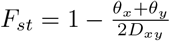^48^, comparing estimated values to the ground truth, calculated by counting substitutions.

#### 2.2.1 DipSkmer provides better *θ* estimates with multiple iterations

Before comparing DipSkmer to other methods, we first compare per-sample and per-pair estimates of *θ*. Recall that we use *k*-mer frequency spectrum to obtain a *θ* for each sample used in coverage calculations (Section 4.4), in addition to the pairwise estimate of *θ* based on Jaccard. In simulated datasets, pairwise Jaccard-based estimates of *θ* are far more accurate than within-sample frequency-spectrum-based estimates, especially with low coverage (Fig. S2). Based on this observation, we propose an iterative approach to estimate *θ* using the within-sample method, estimate coverage, estimate pairwise *θ*, and use this new *θ* to obtain better coverage, iterating this process several times. DipSkmer’s average error reduces from 46% with the within-sample method to 29% error in the third pass of the iterative approach (Fig. S3). With the exception of high coverages (4× and 8×), the third iteration of DipSkmer is more accurate than using within-sample estimates from Equation (7). On empirical data, correlations stay stable across passes (Fig. S4). However, going from one pass to three passes tends to help accuracy in cases where the true *θ* is much lower than the starting *θ*^*\**^ used in the first pass (0.005). While it is not universally the best, the third pass is used in all subsequent DipSkmer results.

#### 2.2.2 DipSkmer *θ* estimates are more accurate than haploid methods

Across most coverages, populations, and parental genomes, DipSkmer provides substantial improvements compared to ReSkmer-ref, Skmer, and Mash (Fig. 2). Overall, DipSkmer has the smallest MAE (25%), which is a substantial improvement compared to ReSkmer (67%), Skmer (86%) and Mash (619%) (Fig. S5). Comparing coverage levels, DipSkmer obtains its highest accuracy at 2× (with 17% MAE) where it provides substantial improvements over ReSkmer-ref (57%), Skmer (81%) and Mash (476%). Only at 8×, the highest simulated coverage, does Skmer achieve comparable accuracy (19% MAE) to DipSkmer. Mash has the best accuracy at 8× coverage when *θ* is ≥ 0.009, but is otherwise inaccurate; it has very high error when coverage is below 4× or when estimating small distances (*θ* < 0.005); even at 8× coverage, it has an error of 190% for these small distances.

**Figure 2:**
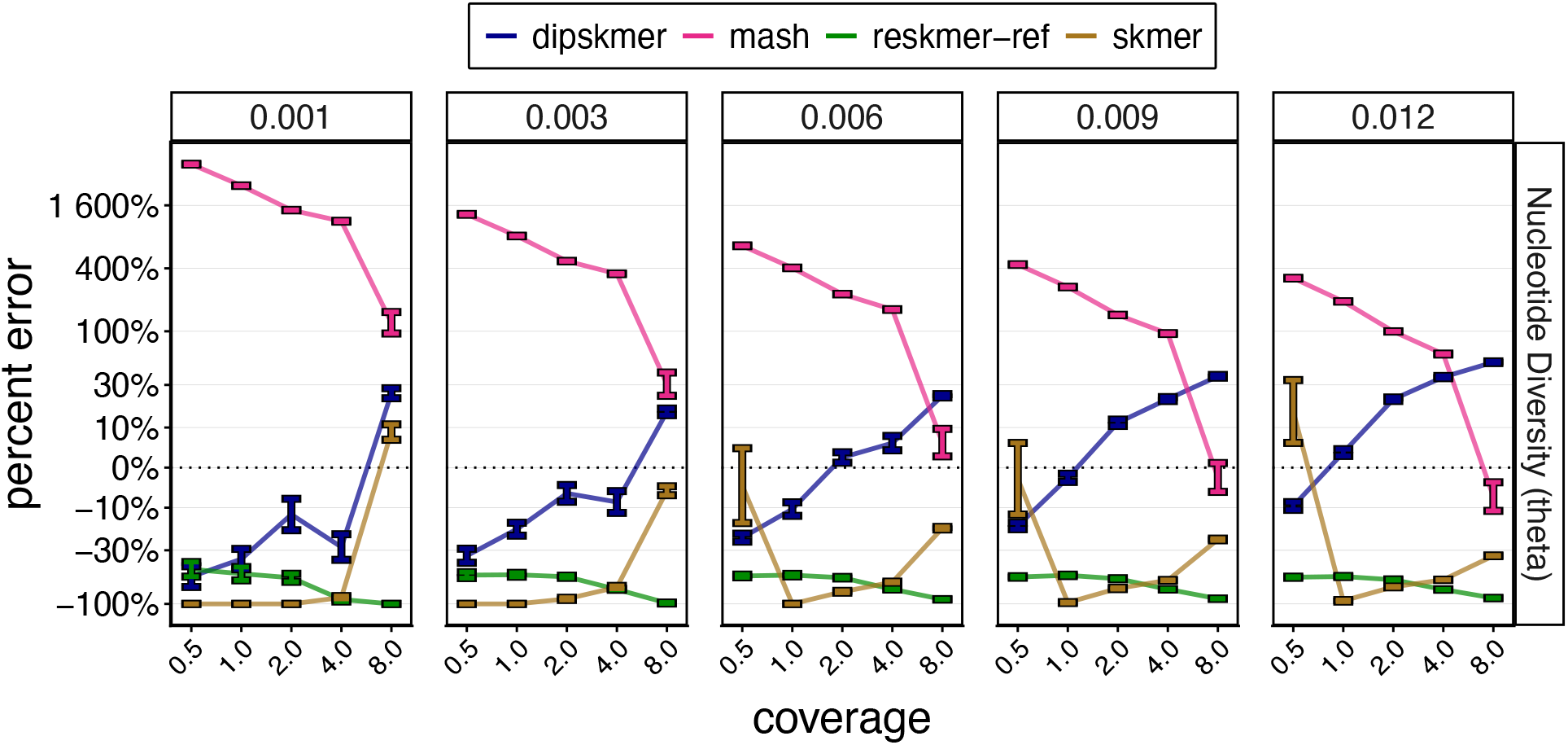
Accuracy of Nucleotide Diversity (*θ*) Estimates on Simulated Data. Percent error 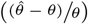 plotted against sequencing coverage. Panels correspond to the true *θ* value for each simulated population size. Bars show the range of error across 5 replicates of three genomes. *Note:* error of -100% means the method estimated 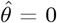.

When comparing methods at very low coverages (0.5× and 1×) DipSkmer is far more accurate than other methods. At these coverages, Skmer consistently outputs an estimate of zero when the simulated *θ* ≤ 0.003. ReSkmer-ref is as accurate as DipSkmer for *θ* ≤ 0.003, but not for higher *θ*. Examining each of the three parental genomes separately (Figs. S5 and S6), we observe that DipSkmer is especially accurate in the least repetitive genome we simulate (leech) with a MAE of 20% (compared to ReSkmer-ref’s 79%, Skmer’s 70%, and Mash’s 626%). DipSkmer’s error for the two more repetitive genomes only increases to 28%, while Skmer error rises to 97%. Among these moderately repetitive genomes (UR>0.7), ReSkmer-ref becomes more accurate as repeat content increases, but remains less accurate than DipSkmer in most situations. However, for extremely repetitive genomes (*UR* ∈ {0.6; 0.5; 0.4}) and low *θ* (*θ* = 0.001 − 0.003), ReSkmer-ref becomes more accurate than DipSkmer (Fig. S7), which often produces 0 distance. Even for these genomes, DipSkmer is competitive with ReSkmer at higher *θ* values, outperforming ReSkmer at low coverages.

We also test how accurately DipSkmer estimates sequencing parameters compared to Skmer and ReSkmer (Fig. S8). In terms of coverage, error rate, and genome length, DipSkmer clearly outperforms Skmer. There is a large improvement in sequencing error estimates at 0.5× and 1× coverage, with more modest gains for higher coverages. The estimated coverage and genome length similarly improve the most at lower coverages but gain very little at higher coverages. ReSkmer-ref understandably outperforms DipSkmer in genome length and coverage estimations, given we are providing an error-free reference genome. In spite of not being reference-based, DipSkmer obtains more accurate sequencing error estimates than ReSkmer-ref when coverage ≥ 2×. Despite these improvements, which are likely due to modeling within-sample *θ* in repeat-spectrum analyses, some overestimation of coverage and thus underestimation of genome length remain, perhaps due to inaccuracies in estimating within-sample *θ* (Fig. S2).

#### 2.2.3 DipSkmer accurately measures population divergence with *D*_*xy*_ **and** *F*_*ST*_

We next measure DipSkmer’s ability to measure divergence between populations in simulated data, starting with the fixation index, Weir and Cockerham *F*_*ST*_ (Fig. 3). DipSkmer and Skmer both overestimate *F*_*ST*_, but DipSkmer has a lower error in most conditions, with some exceptions among high coverage and high divergences (coalescent unit 1 divergence, which often corresponds two different species). Notably, with lower coverage, DipSkmer is much more accurate than Skmer, and its accuracy is highest at around 1–2× coverage. Mash has the opposite error, underestimating *F*_*st*_ in all cases with coverage below 8×. Mash essentially fails to detect differentiation between populations when coverage is below 4×. Conversely, Mash is very accurate when the population differentiation is at the speciation level (1 coalescent unit) and coverage is high (i.e., conditions it is designed for); in these conditions, which strongly break DipSkmer’s assumptions of random coalescence, DipSkmer fails to detect differentiation at 4–8× coverage. Nevertheless, DipSkmer produces accurate estimates at 0.5–2× coverage despite strong model misspecification. We also test two more definitions of *F*_*ST*_, Nei’s and Slatkin’s, and observe similar results (Fig. S9). Repeats play a role in all methods’ *F*_*st*_ estimation accuracy (Fig. S10); DipSkmer achieves its highest accuracy in low repeat genomes and becomes less accurate in higher repeat genomes. However, the relative accuracy of methods remained similar across repeat levels.

**Figure 3:**
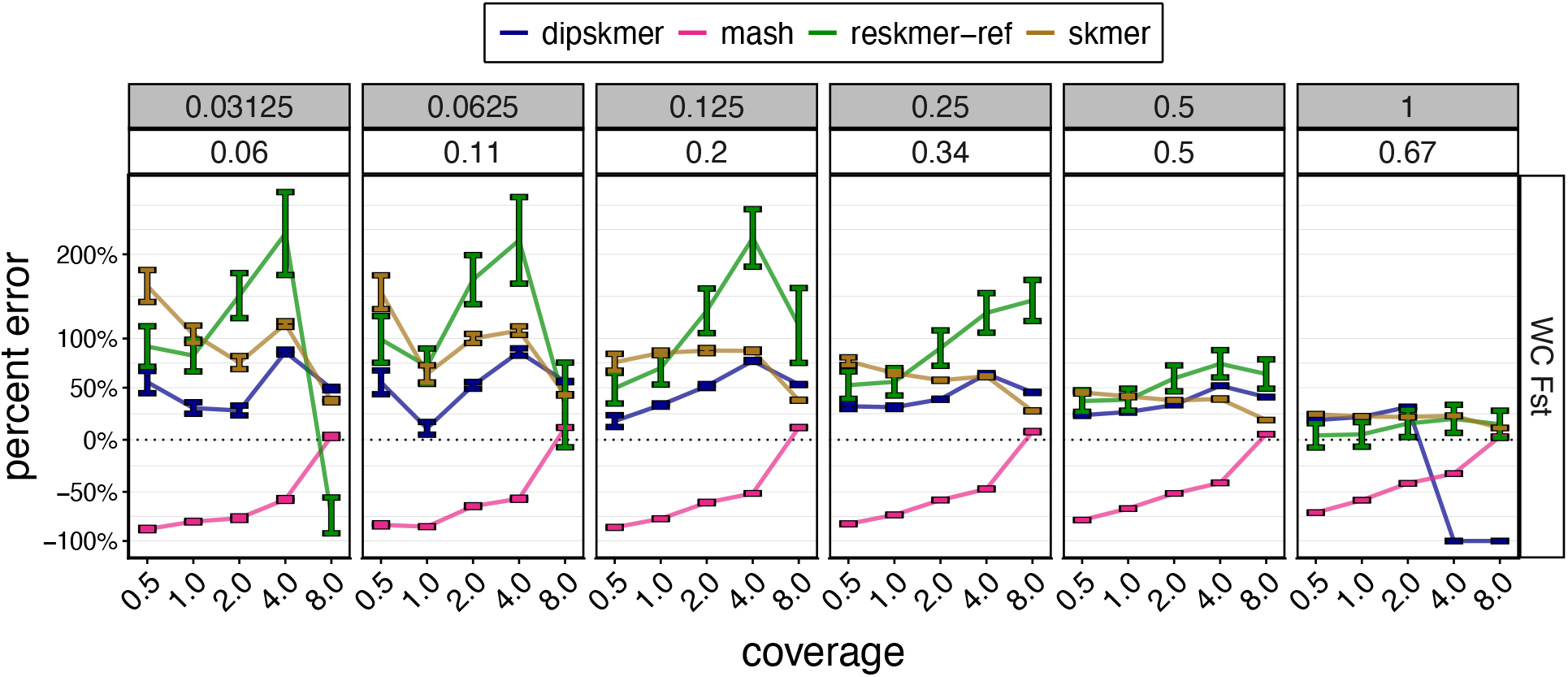
Accuracy of fixation index (*F*_*st*_) on simulated data. We show results for Weir and Cockerham *F*_*st*_ across two simulated populations at a constant population size (*N*_*e*_ = 5 × 10^6^). Panels correspond to different levels of divergence in coalescent units (gray strip) with the corresponding mean true WC *F*_*st*_ shown below.

Another measure of divergence we test is the average distance between samples of different populations, *D*_*xy*_ (Fig. S11). For *D*_*xy*_, DipSkmer is the most accurate method in most conditions, with some exceptions when divergence is very high or the coverage is 8×. Skmer and Mash both perform better than DipSkmer at 8× coverages. ReSkmer-ref has a similar performance to Skmer, but becomes more inaccurate as coverage increases. The *D*_*xy*_ metric clearly shows the impact of population divergence on DipSkmer: As population divergence increases (breaking random coalescence assumptions), the accuracy of *D*_*xy*_ declines steadily, especially above 0.25 coalescent units divergence. In contrast to *F*_*st*_ and *θ*, repetitiveness does not affect *D*_*xy*_ estimation accuracy much. DipSkmer’s accuracy stays around 31%-32% regardless of repeat level (Fig. S11).

### 2.3 Biological Results

To test DipSkmer on empirical data, since true *θ* values are not known, we compare *k*-mer-based methods to previously published^26^ *θ* estimates obtained by ANGSD^21^, which is a widely used method based on read mapping, genotyping, and computing Site Frequency Spectrum (SFS). The dataset is composed of 6 species and 12 populations (Table S2). To get measures of correctness of our nucleotide diversity estimates, we measure the difference between *θ* values and the Pearson correlation of our estimates of *θ* to those produced by ANGSD. We follow a published protocol^27^ for the preparation of reads (described in Methods). We then subsample all genome skims to target coverages *c* ∈ {0.5×, 1×, 2×, 4×, 6×, 8×}.

#### 2.3.1 DipSkmer closely matches ANGSD *θ* estimates in empirical datasets

On the biological dataset, to gauge DipSkmer’s ability to quantify relative changes in nucleotide diversity, we first test how well DipSkmer correlates with ANGSD estimates of *θ* (Fig. 4A). According to the Pearson correlation coefficient (*R*), DipSkmer has the highest correlation averaged across coverages (*R*=0.92), followed closely by Skmer (*R*=0.9) and ReSkmer-noref (*R*=0.9), all of which perform far better than Mash (*R*=0.43). Mash is essentially uncorrelated at lower coverages, starts to have some correlation at 2×, and matches other methods at 6× or higher. DipSkmer, in contrast, has high correlation coefficients at all coverage levels, with minimal change across coverage.

**Figure 4:**
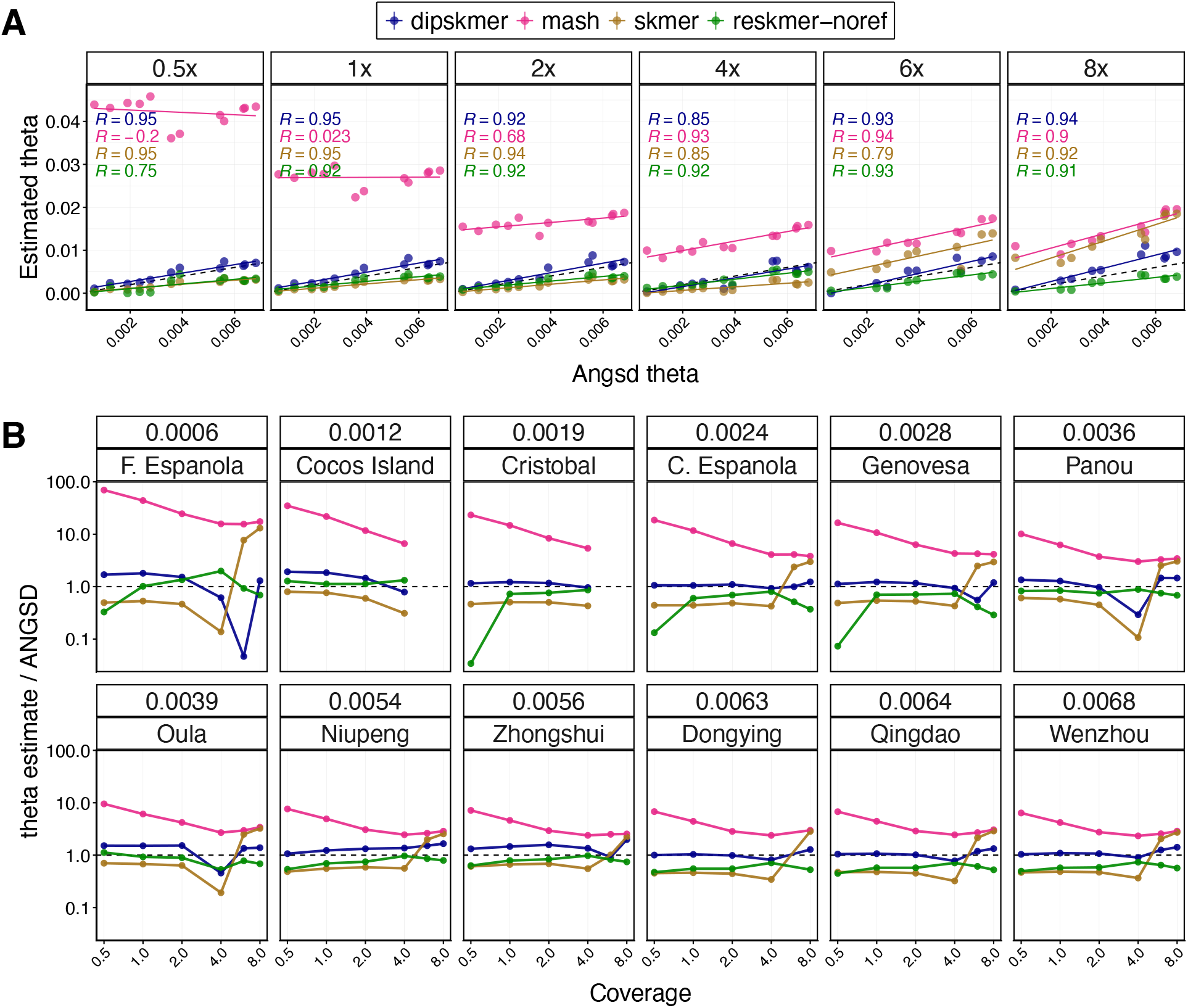
Correlations and Accuracy in Empirical Datasets. **A)** Correlation of DipSkmer, Skmer, and Mash estimates of *θ* (*y*-axis) to ANGSD (*x*-axis). Each panel corresponds to a different level of coverage. Pearson correlation coefficients (R) are found in the top left corner of each panel. **B)** Ratio (*y*-axis) between DipSkmer, Skmer, and Mash *θ* estimates and ANGSD *θ* across different coverages (*x*-axis). Panels represent different populations, with population name and corresponding ANGSD estimate on the top strips.

Beyond correlations, DipSkmer also closely matches ANGSD estimates of *θ*, in contrast to Mash, which systematically overestimates, and Skmer, which systematically underestimates ANGSD (Fig. 4B). ReSkmer-noref is competitive with DipSkmer, except at 0.5× coverage, where ReSkmer underestimates *θ* in some populations (see C. Espanola, Cristobal, and Genovesa). Across all populations and coverage, DipSkmer produces the most similar distances to ANGSD judged by MAE (28%) compared to ReSkmer-noref (41%), Skmer (83%), and Mash (738%); this pattern remains correct for each coverage level (Table S1). Overall, the impact of coverage on DipSkmer is relatively low, with MAE ranging between 22% and 40%, with the peak accuracy achieved at 2× coverage. Meanwhile, ReSkmer-noref achieves its peak accuracy at 4× coverage (30% MAE), where it is nevertheless outperformed by DipSkmer (26%).

For *Sillago Sinica* populations (Dongying, Qingdao, and Wenzhou) we test the similarity of reference-free estimates of *θ* between pairs of samples to those produced by an ANGSD-produced identity by state (IBS) matrix. IBS measures a normalized genetic distance between aligned loci given reference genomes and high coverage samples – which DipSkmer does not require. Again, DipSkmer produces the most similar estimates to ANGSD (Fig. S12), with a mean percent difference of 12% across all coverages. The two next most accurate methods have similar percent error: ReSkmer-noref (30%) and Skmer (43%). There is no substantial change in error between coverages, except for Mash whose error decreases from 687% (0.5×) to 237% (6×). Notably, even though coverage is relatively high, Mash remains highly inaccurate.

#### 2.3.2 DipSkmer best approximates ANGSD estimates of *Sillago sinica* population structure

To test how DipSkmer performs in the presence of population structure, we take three populations from the same species (*Sillago sinica*) and look at two metrics for comparison: *F*_*st*_ and Principal Coordinates Analysis (PCoA). We perform PCoA on identity by state (IBS) matrices produced by ANGSD as well as those produced by reference-free methods to project distances onto Euclidean space, followed by Procrustes to align the projections to enable comparison across methods.

Projections based on DipSkmer distances are remarkably close to those based on ANGSD (Fig. 5A). In fact, in most cases, ANGSD projections are within the confidence intervals for DipSkmer (obtained using subsampling reads, see Methods). Skmer and ReSkmer-ref also capture the broad patterns but show lower similarity in their projections to those of ANGSD, in particular for four outlier Dongying samples. Judged by the alignment of points to ANGSD, as measured by the *t*_0_ similarity score (see Methods), DipSkmer is closest to ANGSD across all coverages and achieves very high similarity (*t*_0_ = 0.821) for 2× – 6× (Fig. 5B). All methods have lower *t*_0_ values at coverage ≤ 1×, reach their peak at 2×, with little change at higher coverages.

**Figure 5:**
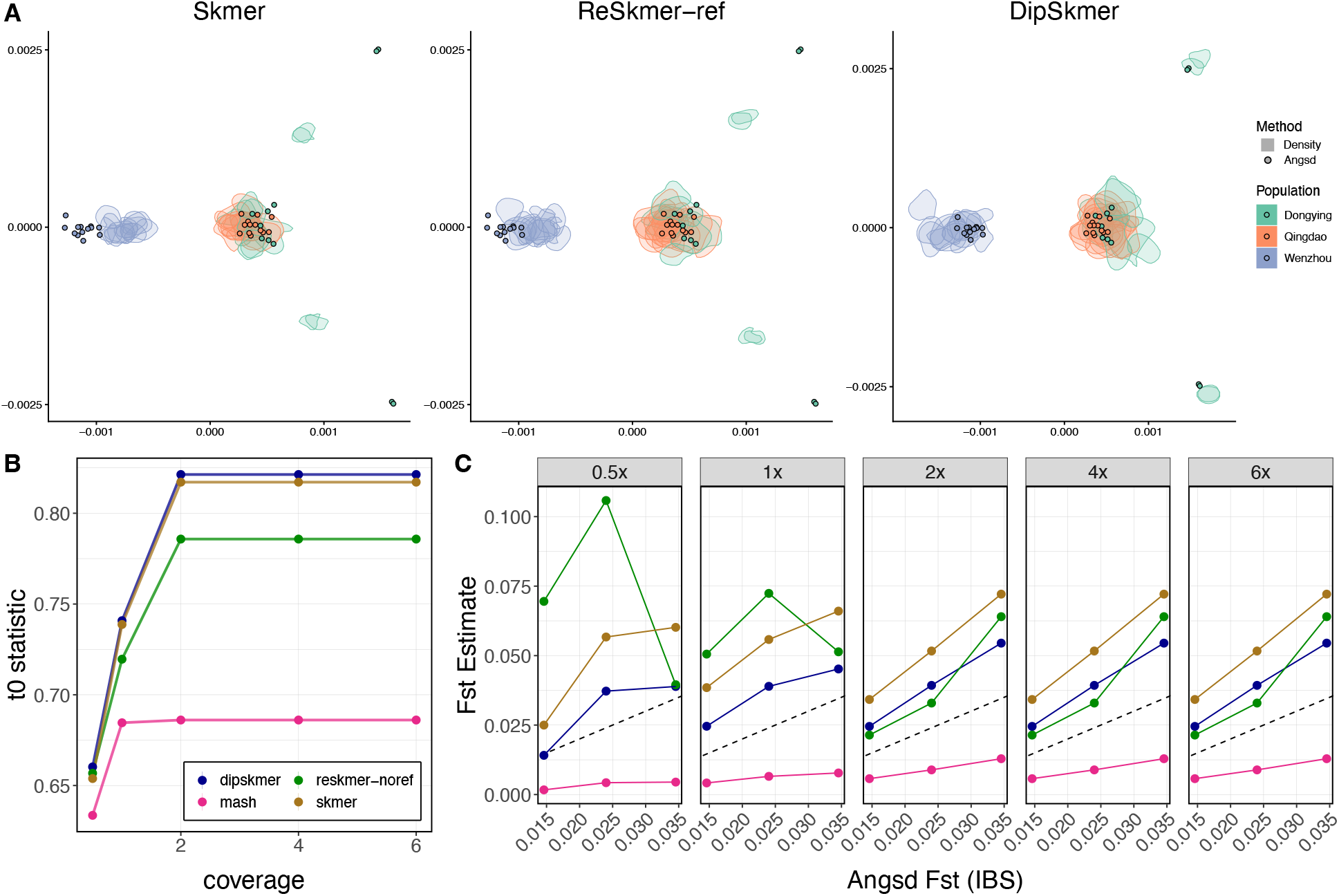
*Sillago sinica* Population Structure: **A)** Similarity between PCoA produced by ANGSD and those produced by three different alignment-free methods. Colors represent three different *Sillago sinica* populations. Points show the coordinates of ANGSD PCoA results and contours show the location of the highest 33% density of data points for each sample using the corresponding method. **B)** Similarity in PCoA structure (*t*_0_) between PCoA produced by ANGSD and those produced by skimming methods. The *t*_0_ statistic ranges from 0 (dissimilar) to 1 (identical). **C)** Correlation plot between reference-free estimates of Weir and Cockerham *F*_*st*_ (*y*-axis) and those produced using the ANGSD IBS matrix (*x-axis*). The dotted line denotes the optimal relationship between estimates.

To test the accuracy of *F*_*st*_ measurements across methods, we use the distances output by reference-free methods to compute Weir and Cockerham (WC) *F*_*st*_ (see Methods) and compare those to WC *F*_*st*_ calculated from ANGSD IBS matrices (Fig. 5C). At 0.5× and 1×, DipSkmer produces the most similar estimates to ANGSD. However, at > 1×, ReSkmer-noref produces slightly more similar estimates to ANGSD for two population pairs, Qingdao-Dongying and Wenzhou-Qingdao, followed by DipSkmer. Finally, we also compare our *F*_*st*_ values to those produced by 2D-SFS analysis with ANGSD (Fig. S13). Here, again, DipSkmer is the most accurate among reference-free methods at low coverages (22% error when coverage < 2×).

## 3 Discussion

Here we presented DipSkmer, a method that models diploidy’s effect on the Jaccard index of *k*-mer sets to produce accurate and robust estimates of nucleotide diversity *θ* from diploid samples. In both simulated and empirical datasets, DipSkmer outperformed other alignment-free, reference-free methods for diploid inputs. Additionally, it was able to estimate measures of population differentiation such as *F*_*st*_ and *D*_*xy*_ with high accuracy. Moreover, DipSkmer is fast, requiring only 5 minute on average on our datasets for each sample (Table S4), which is identical to Skmer, and not much higher than Mash, which needs one minute per sample.

The DipSkmer’s promise for practical biodiversity monitoring lies in providing estimates that highly correlate with the more laborious option of mapping reads to a reference genome (ANGSD). Crucially, since DipSkmer eliminates the need for access to reference genomes, it opens up the possibility for population monitoring using genome skims for less well-studied species. And the fact that it requires low coverage reduces the cost. In fact, DipSkmer was generally more accurate with 2× coverage than 4×, which may seem counterintuitive. We refer to Remark S1 (in supplementary material) for some theoretical insights as to why this may be the case: Around 4× coverage, changing *θ* surprisingly has little to no effect on how many *k*-mers are shared between two samples, reducing sensitivity. In practice, when coverage is 3–5×, we suggest downsampling reads to 2×.

The theory we provide allows estimation of *θ* from a single sample using its *k*-mer repeat spectrum or using two samples by modeling Jaccard. Given the ability to estimate heterozygosity from a single sample, which others have also attempted^19^, one may wonder why the Jaccard-based estimates were necessary. Our empirical results showed that within-sample estimates of *θ* are far less robust, especially with low-coverage data (Fig. S2). In contrast, with correct coalescent modeling, Jaccard distances between pairs of samples provide highly accurate estimates of *θ*. Thus, we advise practitioners to use the pairwise estimates rather than single-sample estimates. Pairwise estimates additionally enable comparing populations (e.g., *D*_*xy*_ and *F*_*st*_), which is not possible with single-sample *k*-mer statistics.

While our theory was based on Wright Fisher (WF) populations with random mating, DipSkmer was empirically robust when applied to individuals from two divergent populations, including at very low coverage and low population divergence. In extreme simulated cases where population separation approached the speciation-level divergences (e.g., around 1 coalescence unit in simulations), the accuracy of DipSkmer distance estimates drops. Thus, the use of this model should clearly be restricted to comparisons within a species or very close species. When populations are highly diverged, the impacts of within-sample heterozygosity become negligible compared to phylogenetic divergence, making other methods such as ReSkmer, Skmer, or Mash more appropriate. Despite our empirical robustness to moderate levels of population structure, future research should explore how differing coalescence dynamics in demographic models with population isolation and migration^31,49^ affect the Jaccard index in theory.

Just as DipSkmer is an extension of Skmer that models heterozygosity, the recent ReSkmer method is an extension that models repeats based on *k*-mer frequency spectrum, which can be computed using Respect^44^. Unlike DipSkmer, ReSkmer (and Respect) ignore the impacts of heterozygosity. One should ideally model both aspects, but analytical equations become intractable. While we leave it to future work to combine the two models, we can empirically ask which aspect has a higher impact – accounting for repeats or heterozygosity. Based on our results, overall, heterozygosity appears to have a greater impact on Jaccard than repeats for population genetic level distances, though this pattern is not universal. When repeat content is extremely high and one has access to accurate reference genomes to compute *k*-mer frequency spectrum, ReSkmer can be more accurate (Fig. S7). Our results motivate future attempts to combine repeat modeling with within-sample heterozygosity. Due to the complexity of the resulting equations, the combined analyses may require moving beyond analytical approaches and adopting machine learning, following some earlier attempts^47^. Such combined modeling may also enable us to move beyond diploid organisms and model polyploidy and mixed-ploidy.

## 4 Methods

### 4.1 Background: Jaccard index for two haploids within a coalescence framework

The known connections between Jaccard and *θ* can be arrived at by analyzing the coalescent on *n* = 2 haploid samples from a WF population. Let each *k*-mer be considered a set of *k* independent characters. For haploids, the expected time to coalesce under the WF model is 𝔼[*t*_2_] = *N*_*e*_, and thus, the expected number of mutations on the lineage is Λ := 2*N*_*e*_*µk* = *θk*. The number of mutations can be considered a Poisson process with parameter Λ; then, *Q* := Pr[no mutation on the lineage] = *e*^*−*Λ^ ≃ (1 − *θ*)^*k*^. Thus, for each position of genomes of length *L*, with probability *Q*, a single *k*-mer is added to both the intersection (*I*) and the union (*U*) of *k*-mer sets. If there is at least a mutation (probability 1 − *Q*), the position adds 0 *k*-mers to the intersection and 2 *k*-mers to the union. Thus, 𝔼 [*I*] = *QL* and 𝔼 [*U*] = *L*(2(1 − *Q*)+ *Q*). Using the (biased) ratio estimator, one can arrive at 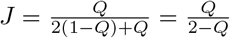, which, solved for *θ* would give 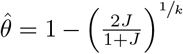.

These calculations fail at low coverage and ignore sequencing errors^43^. Define *L* := length of haploid genome, *N* := number of sampled base pairs, *ℓ* := read length, 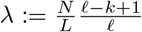 *k*-mer coverage. We assume each *k*-mer has probability *ρ* := (1−*ϵ*)^*k*^ ≃ *e*^*−kϵ*^ of being error-free and let *ξ* := *λρ* be the error-free coverage of *k*-mers. Sarmashghi et al.^43^ showed that *ϵ, λ* can be estimated for each sample from their *k*-mer frequency profile. Assuming random errors, η := Pr[an arbitrary k-mer at a specific location is sampled without error] = 1 − *e*^*−ξ*^. Assuming errors generate unique *k*-mers, ζ := 𝔼[the number of unique *k*-mers generated from a position] = η + *λ* − *ξ*, where η captures the error-free *k*-mers and the other terms the erroneous ones. Sarmashghi et al.^43^ showed that with those estimates at hand for both samples (denoted by subscript), the hamming distance (thus, *θ*) can be estimated using

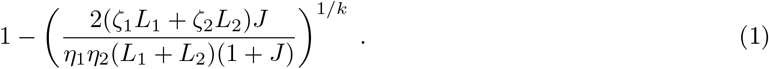

### 4.2 Extending to two diploid individuals (*n* = 4)

We now turn to the diploid case, where we assume *n* = 4 haploids are sampled from a randomly mating population, so that coalescence between parental and maternal alleles of a sample is identically distributed to alleles from different individuals. We start by analyzing genomic *k*-mers (i.e., all *k*-mers are sampled without error), then model the impacts of low coverage and sequencing errors. Recall that for diploids, *θ* := 4*N*_*e*_*µ*.

Consider the coalescence history of each locus, which can take one of two shapes (Fig. 6a), with the unbalanced case twice as likely as the balanced. Under Kingman’s coalescence^20^, for each time epochs *t*_*i*_ (*i* ∈ {2, 3, 4}) with i lineages, 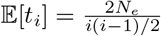. The expected total branch length is 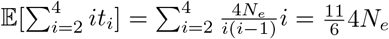. Under the infinite sites assumption, the number of substitutions per *k*-mer falling on these genealogies is Poisson distributed with parameter 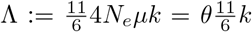 and *Q* := Pr[no mutation in lineage of 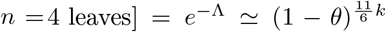 (compare to haploids). With probability *P* := 1 − *Q*, we have one or more substitutions in a *k*-mer. We only analyze the possibility of zero or one substitutions, essentially assuming that in the unlikely event that multiple substitutions occur, the ratio of the expected number of shared *k*-mers to all *k*-mers is well approximated by derivations for one substitution. Conditioned on having exactly one substitution, *p*_*i*_ := Pr[substitution falls on a specific lineage of epoch 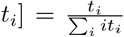, and thus, *p*_2_ =^3^*/*^11^, *p*_3_ =^1^*/*^11^, *p*_4_ =^1^*/*^22^.

**Figure 6:**
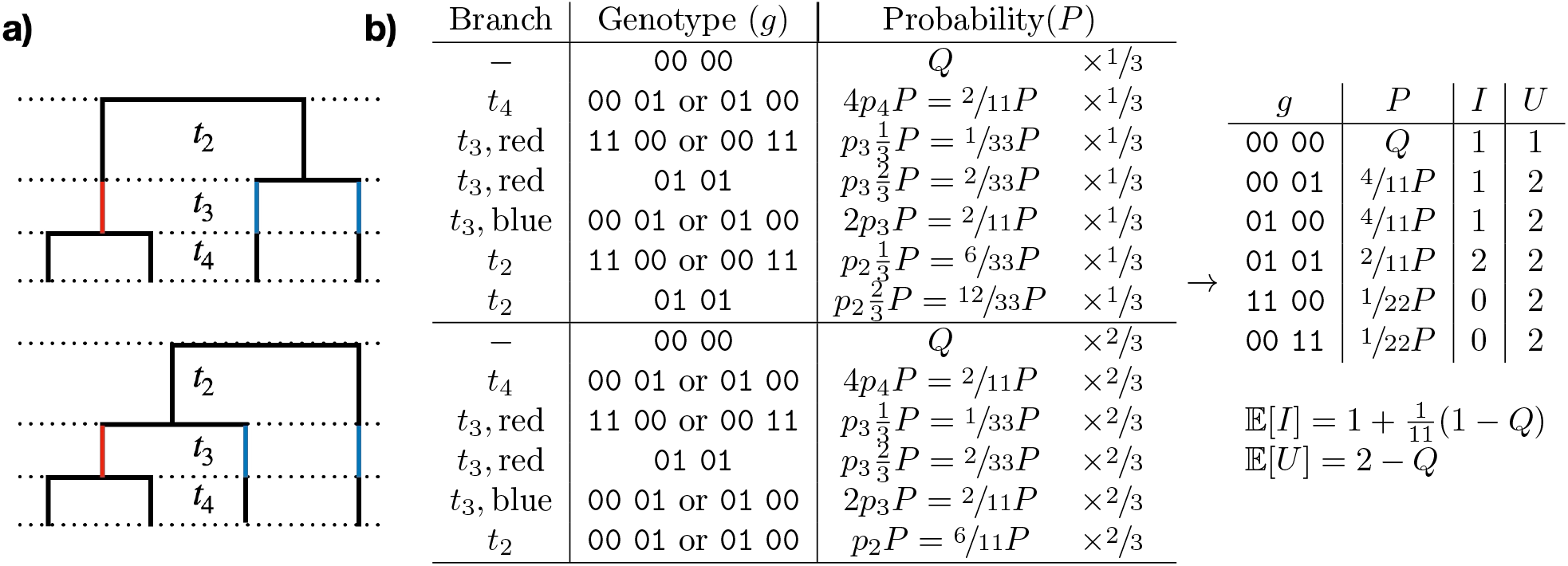
a) Possible coalescence topology shapes. Assignment of alleles to leaves is uniformly random. The epochs *t*_2_ … *t*_4_ are marked. Note that in *t*_3_, the red branch can coalesce with either blue ones to give the unbalanced tree, and thus, the probability of the unbalanced tree is 2*/*3 versus 1*/*3 for balanced. b) If a substitution occurs (probability *P* = 1 *− Q*), depending on which branch it falls on, it leads to different genotypes. Each genotype contributes 0–2 *k*-mers to the intersection (*I*) and 1 or 2 *k*-mers to the union *U* of the *k*-mer sets obtained from two diploid genomes.

We assign a random permutation of haplotypes to the leaves of the topologies and analyze the expected number of *k*-mers in the intersection (*I*) and union (*U*) for a single genomic position. Figure 6B summarizes analyses of both topologies. With probability *Q*, no substitutions occur, which leads to *I* = 1, *U* = 1. If any mutation occurs, *U* = 2. If it falls on any lineage in *t*_4_, the genotypes of the two individuals is some permutation of 00 01, which means, *I* = 1. If the mutation falls on the first lineage in *t*_3_, the two lineages carrying the mutation may belong to the same individual (this happens in 8 out of 24 =^1^*/*^3^ of all permutations), resulting in *I* = 0, or they may belong to different individuals, leading to *I* = 2. If the mutation falls on other lineages of *t*_3_, then *I* = 1. The *t*_2_ epoch depends on the topology, where the balanced tree has the same scenarios as *t*_3_, leading to *I* = 0 or (twice as likely) *I* = 2. For unbalanced, regardless of the permutation, both samples will have one of each allele, leading to *I* = 2. Listing all scenarios, we conclude: 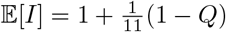 and 𝔼[*U*] = 2 − *Q*. Using the same method of ratios as Mash/Skmer and solving for *Q*, we obtain:

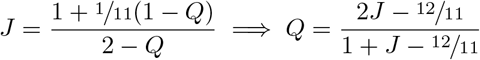

which adds −^12^*/*^11^ to numerator and denominator of the haploid equation. Setting 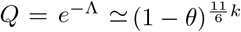 gives:

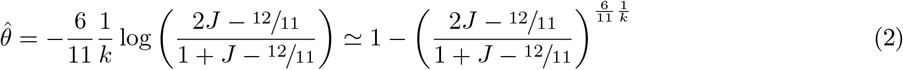

This diploid estimate of 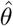 gives a higher value than the haploid estimate (Fig. S1A) and is only defined for J >^6^/^11^. To see why, note that under the infinitive sites assumption (which breaks at higher distances), two samples with high *θ* would still have substantial similarity because the two alleles of each individual are sampled from the same pool of alleles available in the population. As shown in Section SA.2.2, a haploid estimate based on Jaccard can be transformed to the diploid estimate using a non-linear equation.

### 4.3 Modeling low coverage and errors

We next study Jaccard for genome skims where some *k*-mers are not covered and others are erroneous. Recall definitions of *L, N, ℓ, ρ, ξ* from Section 4.1. We henceforth update the definition of 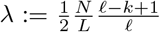 to be the *haploid k*-mer coverage for a diploid skim, and note that this is half of what we defined before. We aim to model the fact that due to sequencing errors and limited sequencing coverage, a *k*-mer could be missing and/or sampled erroneously in one or both samples, thereby reducing the intersection and potentially increasing the union. Mash deals with errors by requiring each *k*-mer to be observed at least *t* times, noting that for a threshold *t* sufficiently large compared to *λ*, a *k*-mer sampled *t* times is likely to be error-free. This approach only works for high coverage, since for low coverage, very few *k*-mers will survive this filter. Our approach is to select a *t* ≥ 1 based on our estimate of coverage. In practice, we set *t* = ⌊ ^*ξ*^/^2^ ⌋ +1 by default, allowing each sample *j* to have its own *t*_*j*_. We assume that no erroneous *k*-mer will match any alleles in the genome. Let

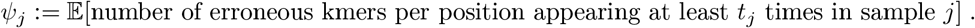

Thus, each position in expectation adds *ψ*_1_ + *ψ*_2_ erroneous *k*-mers to the union and none to the intersection. Since errors are mostly random, with our choice of *t*_*j*_, it is safe to assume *ψ*_*j*_ = 0 for *t*_*j*_ *>* 1 and *ψ*_*j*_ = 2*λ*(1 − *ρ*) for *t*_*j*_ = 1. We now focus on the error-free *k*-mers, which may be homozygous or heterozygous. Let

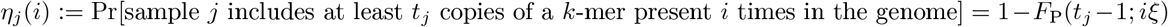

where *F*_*P*_ is the CDF of Poisson, which can be calculated using the regularized incomplete gamma function. Since a *k*-mer may be homozygous or heterozygous (effectively covered twice), we are interested in *i* ∈ {1, 2}. For each location, and for each genotype (Fig. 6), we can compute their expected contribution to *U* and *I*, as listed below, from which 𝔼[*I*] = **P** *·* **I** and 𝔼[*U*] = **P** *·* **U** + *ψ*_1_ + *ψ*_2_ can be calculated (see Eqs. (S5) and (S6)).

| Genotype $g$ | $\mathbf{P}$ | $\mathbf{I} = \mathbb{E}[I\vert g]$ | $\mathbf{U} = \mathbb{E}[U\vert g, \text{error-free}]$ |
| --- | --- | --- | --- |
| 00 00 | $Q$ | $\eta_1(2)\eta_2(2)$ | $\eta_1(2) + \eta_2(2) - \eta_1(2)\eta_2(2)$ |
| 00 01 | $\frac{4P}{11}$ | $\eta_1(2)\eta_2(1)$ | $\eta_1(2) + \eta_2(1) - \eta_1(2)\eta_2(1) + \eta_2(1)$ |
| 10 00 | $\frac{4P}{11}$ | $\eta_1(1)\eta_2(2)$ | $\eta_1(1) + \eta_2(2) - \eta_1(1)\eta_2(2) + \eta_1(1)$ |
| 01 01 | $\frac{2P}{11}$ | $2\eta_1(1)\eta_2(1)$ | $2(\eta_1(1) + \eta_2(1) - \eta_1(1)\eta_2(1))$ |
| 11 00 or 00 11 | $\frac{P}{11}$ | 0 | $\eta_1(2) + \eta_2(2)$ |

For example, for the genotype 00 01, *I* = 1 if and only if the *k*-mer is sampled at least *t*_1_ times from the first skim where it is homozygous, and *t*_2_ from the second skim, where it only appears in one chromosome; otherwise, *I* = 0. Therefore, 𝔼[*I*|00 01] = η_1_(2)η_2_(1). For union *U*, we have two distinct *k*-mers. Allele 0 is sampled at least *t*_1_ and *t*_2_ times without error with probabilities η_1_(2) and η_2_(1) in each individual and η_1_(2)η_2_(1) in both, and allele 1 with probability η_2_(1) in the second individual. Thus, 𝔼[*U* |00 01] = η_1_(2) + η_2_(1) − η_1_(2)η_2_(1) + η_2_(1). Other scenarios are similar. Replacing 𝔼 [*I*] and 𝔼 [*U*] from Equations (S5) and (S6) in 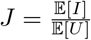 and solving for Q (ratio estimator) results in:

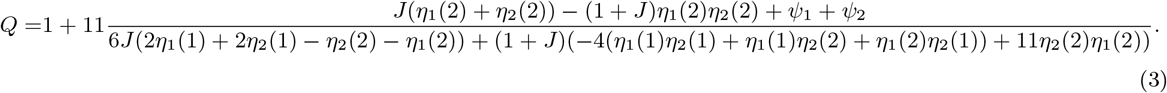

### 4.4 Parameter estimation for diploid skims

To compute Eq. (3), we need to estimate parameters *λ, ϵ* which immediately give us *ξ*, η, *ψ*. Following Skmer, we compute these from the *k*-mer repeat spectrum of each sample, but we model heterozygosity. Let *M*_*i*_ denote the number of *k*-mers observed exactly *i* times and assume all errors fall in *M*_1_. Let

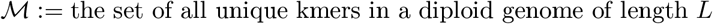

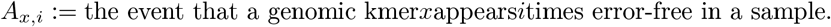

Let *M* := |*M*| and note *M* ≥ *L* and 𝔼[*M*] = *L*(2 − (1 − *θ*)^*k*^) (Section SA.1). Clearly, we need to estimate *θ* for each sample in order to estimate genome length and coverage. Let As shown in Eq. (S1), for i > 1,

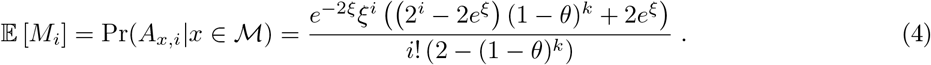

A family of equations can be obtained by taking the ratio of two consecutive instances of Eq. (4) for *i* ≥ 2:

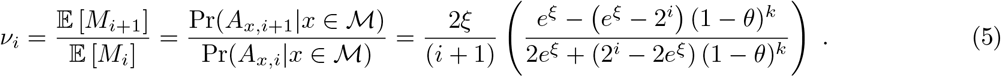

Using *ν*_*i*+1_ and *ν*_*i*_ enables us to completely eliminate *θ* (see Eq. (S3)) and allows us to arrive at an estimate of *ξ*:

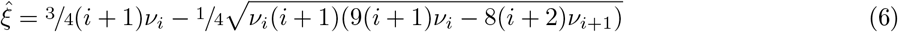

With 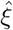 estimated, we then use Eq. (5) to estimate *θ* directly. It can be confirmed that

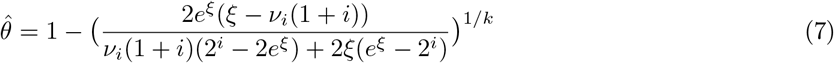

satisfies (5), giving us an estimator for *θ*. While any *i* can be used, we follow Skmer^43^ and set it to arg 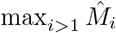 (i.e., the first mode of the skim’s *k*-mer repeat spectrum).

Note that when 9(*i* + 1)*ν*_*i*_ *<* 8(i + 2)*ν*_*i*+1_, Eq. (6) has no solutions, and a different strategy must be used. Let us assume a *θ*^*\**^ is given. Then, we can use Eq. (5) directly to estimate *ξ* by numerically optimizing arg 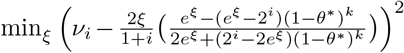. For *θ*^*\**^ we can start with a fixed value (default: *θ*^*\**^ = 0.005). When multiple individuals are available, we can then use an iterative approach to update *θ*^*\**^. In each pass *r*, we use the pairwise sample distances from the previous round (i.e., *θ*_*r−*1_, as described in Section 4.3) as the new *θ*^*\**^ to recompute coverage and error, which we then use to obtain a new pairwise distance *θ*_*r*_. We continue for a fixed number of passes (default: three), and empirically compare this method to using Equation (7).

Regardless of which method is used to compute 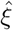, we next need to decompose it into *λ* and *ρ* = (1 − *ϵ*)^*k*^. To do so, we cannot use the family of equations given by (5) because they are only a function of *ξ*. Instead, we need to examine error-prone *k*-mers with *i* = 1. Note that 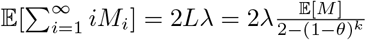.

Then, since all erroneous *k*-mers are assumed to be in *M*_1_, as shown in Eq. (S4), we can drive:

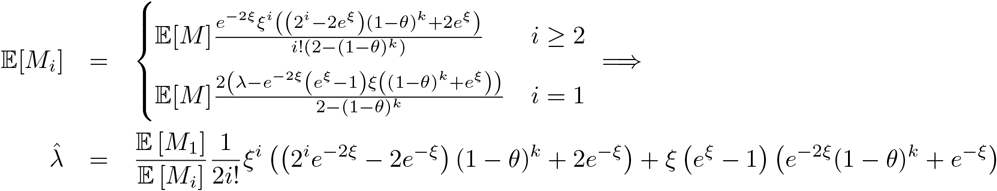

for any i > 1. We can then estimate the error rate from 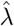 and 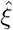 as 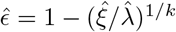.

### 4.5 Experimental setup

#### Simulations

To test the accuracy of DipSkmer, we used the coalescent simulator msprime^3^ to generate four haploid chromosomes with a range of *θ* values. To achieve this, we set rates of mutation (*µ*) and recombination (*r*) to 0.3 × 10^*−*8^ (which is reasonable^32^), and vary effective population sizes (*N*_*e*_ ∈ {8.3 × 10^4^, 2.5 × 10^5^, 5 × 10^5^, 7.5 × 10^5^, 1 × 10^6^}), which lead to *θ* = 4*N*_*e*_*µ* to be 0.001, 0.003, 0.006, 0.009, or 0.012, resp. We repeat the process 5 times for three genome sizes (15 replicates in total). For each replicate, we divide the four chromosomes into two diplid individuals at random. To test whether DipSkmer can also estimate the divergence between populations, we perform separate msprime simulations with a demography where an initial parent population (*N*_*e*_ = 5 × 10^5^) splits into two child populations with no change in *N*_*e*_ and no migration. Two diploid samples are generated for each of the child populations (eight chromosomes in total). To test different levels of divergence, we adjust the number of generations (*τ*) back in time when the populations split (*τ* ∈ {6.25 × 10^4^, 1.25 × 10^5^, 2.5 × 10^5^, 5 × 10^5^, 1 × 10^6^}). These lead to coalescent unit (*τ/*4*N*_*e*_) divergences of {0.03125, 0.0625, 0.125, 0.25, 1}. Note that 1 coalescent unit is high and corresponds to the phylogenetic (rather than population genetic) scale; it is included to test the limits of the methods. Also note that the isolation between populations breaks the random mating assumptions of DipSkmer, and thus, this experiment is meant to test the impacts of model violations.

We generate genomes and genome skims for each simulation. To make these realistic in terms of *k*-mer repeat spectra, we replace the parental genome (root) with one of three assemblies with varying *k*-mer repetitiveness (Table S3). We impose the genotypes (i.e., VCFs) generated by msprime onto the corresponding sites of the genome, selecting among alternative nucleotides at random and leaving the unmutated sites unchanged. For each resulting genome, we use ART^15^ to simulate Illumina short reads (Phred score = 25, *ℓ* = 150) at coverage *c* ∈ {0.5×, 1×, 2×, 4×, 8×}, noting that each of the two diploid chromosomes has a coverage of ^*c*^/^2^. We use the resulting reads as input to DipSkmer, with 1 – 3 passes of *θ*^*\**^ calculations, to DipSkmer with Eq. (5) for coverage, and alternative Jaccard-based methods, Mash^34^ and Skmer^43^. Variants of DipSkmer differ in the calculation of within-sample *θ* used to estimate coverage; however, the accuracy is always measured based on the *θ* obtained from pairs of samples. We measure error as 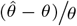, averaged over all replicates, where 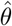 is the estimated pairwise distance and *θ* is the true pairwise distance. We also report mean absolute error (MAE), defined as 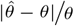. We measure the true distance as the percentage of substituted positions in the simulated genome. For population differentiation experiments, we estimate *D*_*xy*_ (average heterozygosity between pairs of samples taken from different populations) and Weir and Cockerham’s 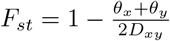, comparing estimated values to the ground truth, calculated based on simulated genomes.

#### Biological data

To test DipSkmer on empirical data, since true *θ* values are not known, we compare DipSkmer, Skmer, and Mash to previously published^26^ *θ* estimates obtained by ANGSD^21^, which is a widely used method based on read mapping, genotyping, and computing Site Frequency Spectrum (SFS). We analyze a dataset^26^ consisting of 12 total populations across six different species with a range of *θ* between 0.0006 − 0.0068 (Table S2). Beyond the difference between *θ* values, we also measure the Pearson correlation of our estimates of *θ* to those produced by ANGSD. To reduce the impacts of contamination on alignment-free methods^38^, we utilize a published protocol^27^, which includes the removal of adapters and deduplication of reads using BBtools^6^. Bacterial and archaeal reads are then removed using CONSULT-II^54^, and human reads are filtered out using Kraken2^50^. To test methods with low-coverage skims, we use Respect^44^ to estimate genome size, then subsample the coverage of all genome skims to target coverages *c* ∈ {0.5×, 1×, 2×, 4×, 6×, 8×}. Note that two populations (Cocos island and Cristobal) were not sequenced deep enough to reach 8× coverage and are absent from high coverage Pearson correlations.

To benchmark DipSkmer’s ability to measure population structure in an empirical dataset, we analyze a short-read WGS dataset of three *Sillago sinica* populations (Wenzhou, Qingdao, Dongying)^53^. To prepare the sequencing reads for ANGSD^21^, we first performed adapter and quality trimming using BBtools^6^ and used BWA-MEM^23^ to map reads to a publicly available reference assembly^51^. We then use ANGSD to obtain an IBS (identity by state) matrix, run a principle coordinate analysis (PCoA) on it, and analyze the first two components. To judge the ability of reference-free methods to obtain population structure, we first used Skmer’s built-in subsampling approach^39^ to generate 100 replicates of reads subsampled from 4× genome skims. We then use Skmer, ReSkmer, and DipSkmer, to construct distance matrices based on those subsamples. Next, we use the Procrustes method to align each of these distance matrices to ANGSD’s PCoA and summarize the results by plotting polygons in regions with 33% highest density of points per sample (representing confidence intervals). To quantify the similarity between ANGSD and other methods at a range of coverages, we use the vegan package^33^ in R to perform a symmetric Procrustes analysis repeatedly (a statistical test referred to as “protest”) to obtain *t*_0_, a statistic derived from the sum of squares (*t*_0_ = 1 − *ss*) between rotated matrices. This tells us how similar projections from reference-free methods are to those produced by ANGSD; *t*_0_ ranges from 0 (dissimilar) to 1 (identical).

To get *F*_*st*_ values with the reference-free methods, we take the distance matrices output by each method and compute Weir and Cockerham’s (WC) definition of *F_st_*^48^ 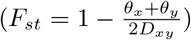. To get *F*_*st*_ with ANGSD, we use two different approaches. First, we take the distances from the IBS matrix to compute WC *F*_*st*_ (referred to as “ANGSD *F*_*st*_ (IBS)”). Second, we use ANGSD and realSFS to generate the 2D Site Frequency Spectra (2D SFS) between all pairs of populations and obtain the unweighted Reynold’s *F*_*st*_^42^.

## 5 Acknowledgements

## 5.1 Contributions

- **Siavash Mirarab:** Conceptualization, Formal Analysis, Funding Aquisition, Supervision, Methodology, Project Administration, Writing - Original Draft Preparation, Writing - Review and Editing, Resources
- **Vineet Bafna:** Conceptualization, Formal Analysis, Funding Aquisition, Supervision, Methodology, Project Administration, Writing - Original Draft Preparation, Writing - Review and Editing
- **Homere J Alves Monteiro:** Investigation, Writing - Review and Editing
- **Eduardo Charvel:** Conceptualization, Investigation, Validation, Visualization, Software, Writing - Original Draft Preparation, Writing - Review and Editing, Formal Analysis, Methodology, Data Curation

## Acknowledgments

We thank Tom Gilbert and Felix Grewe for their conversations and insight. We also Deevanshu Goyal and Adam Bennett for their work in streamlining biological data processing with Skmer. We thank Daira Melendez with her help with empirical datasets. Finally, we thank Ian Kaufman for his expertise and help with HPC systems. This work used Expanse at San Diego Supercomputing Center through allocation ASC150046 from the Advanced Cyberinfrastructure Coordination Ecosystem: Services & Support (ACCESS) program, which is supported by U.S. National Science Foundation grants #2138259, #2138286, #2138307, #2137603, and #2138296.

## References

[1] Daniel N. Baker and Ben Langmead. Dashing: fast and accurate genomic distances with HyperLogLog. Genome Biology, 20(1):265, December 2019. ISSN 1474-760X. doi: 10.1186/s13059-019-1875-0. URL https://genomebiology.biomedcentral.com/articles/10.1186/s13059-019-1875-0.

[2] Pradeepa C. G. Bandaranayake, Nathasha Naranpanawa, C. H. W. M. R. Bhagya Chandrasekara, Hiruna Samarakoon, S. Lokuge, S. Jayasundara, Asitha U. Bandaranayake, D. K. N. G. Pushpakumara, and D. Siril A. Wijesundara. Chloroplast genome, nuclear ITS regions, mitogenome regions, and Skmer analysis resolved the genetic relationship among Cinnamomum species in Sri Lanka. PLOS ONE, 18(9):e0291763, September 2023. ISSN 1932-6203. doi: 10.1371/journal.pone.0291763. URL https://dx.plos.org/10.1371/journal.pone.0291763.

[3] Franz Baumdicker, Gertjan Bisschop, Daniel Goldstein, Graham Gower, Aaron P Ragsdale, Georgia Tsambos, Sha Zhu, Bjarki Eldon, E Castedo Ellerman, Jared G Galloway, et al. Effcient ancestry and mutation simulation with msprime 1.0. Genetics, 220(3):iyab229, 2022.

[4] Kristine Bohmann, Siavash Mirarab, Vineet Bafna, and M. Thomas P. Gilbert. Beyond DNA barcoding: The unrealized potential of genome skim data in sample identification. Molecular Ecology, 29(14): 2521–2534, July 2020. ISSN 0962-1083. doi: 10.1111/mec.15507. URL https://onlinelibrary.wiley.com/doi/abs/10.1111/mec.15507.

[5] A.Z. Broder. On the resemblance and containment of documents. In Proceedings. Compression and Complexity of SEQUENCES 1997 (Cat. No.97TB100171), pages 21–29. IEEE Comput. Soc. ISBN 0-8186-8132-2. doi: 10.1109/SEQUEN.1997.666900. URL http://ieeexplore.ieee.org/document/666900/.

[6] Brian Bushnell, Jonathan Rood, and Esther Singer. BBMerge – Accurate paired shotgun read merging via overlap. PLOS ONE, 12(10):1–15, 2017. doi: 10.1371/journal.pone.0185056. URL https://dx.plos.org/10.1371/journal.pone.0185056.

[7] Selahattin Baris Cay, Yusuf Ulas Cinar, Selim Can Kuralay, Behcet Inal, Gokmen Zararsiz, Almila Ciftci, Rachel Mollman, Onur Obut, Vahap Eldem, Yakup Bakir, and Osman Erol. Genome skimming approach reveals the gene arrangements in the chloroplast genomes of the highly endangered Crocus L. species: Crocus istanbulensis (B.Mathew) Rukšāns. PLOS ONE, 17(6):e0269747, June 2022. ISSN 1932-6203. doi: 10.1371/journal.pone.0269747. URL https://dx.plos.org/10.1371/journal.pone.0269747.

[8] Gerardo Ceballos, Paul R. Ehrlich, Anthony D. Barnosky, Andrés García, Robert M. Pringle, and Todd M. Palmer. Accelerated modern human–induced species losses: Entering the sixth mass extinction. Science Advances, 1(5):e1400253, June 2015. ISSN 2375-2548. doi: 10.1126/sciadv.1400253. URL https://www.science.org/doi/10.1126/sciadv.1400253.

[9] José Cerca Patricia Jaramillo Díaz, Clément Goubert, Heidi Yang, Vanessa C. Bieker, Mario Fernández-Mazuecos, Pablo Vargas, Rowan Schley, Siyu Li, Juan Ernesto Guevara-Andino, Bent Petersen, Gitte Petersen, Neelima R. Sinha, Lene R. Nielsen, James H. Leebens-Mack, Gonzalo Rivas-Torres, Loren H. Rieseberg, and Michael D. Martin. Genomic stability in the Galápagos Scalesia adaptive radiation: Consistent transposable element accumulation despite hybridization and ecological niche shifts, October 2024. URL http://biorxiv.org/lookup/doi/10.1101/2024.09.30.614436.

[10] Tyler K. Chafin, Binod Regmi, Marlis R. Douglas, David R. Edds, Karma Wangchuk, Sonam Dorji, Pema Norbu, Sangay Norbu, Changlu Changlu, Gopal Prasad Khanal, Singye Tshering, and Michael E. Douglas. Parallel introgression, not recurrent emergence, explains apparent elevational ecotypes of polyploid Himalayan snowtrout. Royal Society Open Science, 8(10):210727, October 2021. ISSN 2054-5703. doi: 10.1098/rsos.210727. URL https://royalsocietypublishing.org/doi/10.1098/rsos.210727.

[11] Eduardo Charvel, Isaac Thomas, Homère J. Alves Monteiro, Shahab Sarmashghi, Glenn Dunshea, Vineet Bafna, and Siavash Mirarab. ReSkmer: modeling repeats allows k-mer-based alignment-free methods to calculate population genomic distances. Genome Biology, May 2026. ISSN 1474-760X. doi: 10.1186/s13059-026-04108-9. URL https://link.springer.com/10.1186/s13059-026-04108-9.

[12] Eric Coissac, Peter M. Hollingsworth, Sébastien Lavergne, and Pierre Taberlet. From barcodes to genomes: extending the concept of DNA barcoding. Molecular Ecology, 25(7):1423–1428, April 2016. ISSN 09621083. doi: 10.1111/mec.13549. URL http://doi.wiley.com/10.1111/mec.13549. ISBN: 0962-1083.

[13] Han-Ning Duan, Yin-Zi Jiang, Jun-Bo Yang, Jie Cai, Jian-Li Zhao, Lu Li, and Xiang-Qin Yu. Skmer approach improves species discrimination in taxonomically problematic genus Schima (Theaceae). Plant Diversity, 46(6):713–722, November 2024. ISSN 24682659. doi: 10.1016/j.pld.2024.06.003. URL https://linkinghub.elsevier.com/retrieve/pii/S2468265924000945.

[14] Huan Fan, Anthony R. Ives, Yann Surget-Groba, and Charles H. Cannon. An assembly and alignment-free method of phylogeny reconstruction from next-generation sequencing data. BMC Genomics, 16(1):522, December 2015. ISSN 1471-2164. doi: 10.1186/s12864-015-1647-5. URL http://www.biomedcentral.com/1471-2164/16/522.

[15] Weichun Huang, Leping Li, Jason R. Myers, and Gabor T. Marth. ART: a next-generation sequencing read simulator. Bioinformatics, 28(4):593–594, 2 2012. ISSN 1460-2059. doi: 10.1093/bioinformatics/btr708. URL https://academic.oup.com/bioinformatics/article-lookup/doi/10.1093/bioinformatics/btr708.

[16] A. Randall Hughes, Brian D. Inouye, Marc T. J. Johnson, Nora Underwood, and Mark Vellend. Ecological consequences of genetic diversity. Ecology Letters, 11(6):609–623, June 2008. ISSN 1461-023X, 1461-0248. doi: 10.1111/j.1461-0248.2008.01179.x. URL https://onlinelibrary.wiley.com/doi/10.1111/j.1461-0248.2008.01179.x.

[17] IPBES. Global assessment report on biodiversity and ecosystem services of the intergovernmental Science-Policy platform on biodiversity and ecosystem services, 2019.

[18] Chirag Jain, Luis M Rodriguez-R, Adam M Phillippy, Konstantinos T Konstantinidis, and Srinivas Aluru. High throughput ANI analysis of 90K prokaryotic genomes reveals clear species boundaries. Nature Communications, 9(1):5114, December 2018. ISSN 2041-1723. doi: 10.1038/s41467-018-07641-9. URL http://www.nature.com/articles/s41467-018-07641-9. arXiv: http://dx.doi.org/10.1101/225342.

[19] Katharine M. Jenike, Lucía Campos-Domínguez, Marilou Boddé, José Cerca Christina N. Hodson, Michael C. Schatz, and Kamil S. Jaron. k -mer approaches for biodiversity genomics. Genome Research, page genome;gr.279452.124v1, January 2025. ISSN 1088-9051, 1549-5469. doi: 10.1101/gr.279452.124. URL http://genome.cshlp.org/lookup/doi/10.1101/gr.279452.124.

[20] J F C Kingman. On the genealogy of large populations. Journal of Applied Probability, 19(1982):27–43, 1982. ISSN 00219002. URL http://www.jstor.org/stable/3213548.

[21] Thorfinn Sand Korneliussen, Anders Albrechtsen, and Rasmus Nielsen. ANGSD: Analysis of next generation sequencing data. BMC Bioinformatics, 15(1):356, November 2014.

[22] David Koslicki and Hooman Zabeti. Improving MinHash via the containment index with applications to metagenomic analysis. Applied Mathematics and Computation, 354:206–215, August 2019. ISSN 00963003. doi: 10.1016/j.amc.2019.02.018. URL https://linkinghub.elsevier.com/retrieve/pii/S009630031930116X.

[23] Heng Li. Aligning sequence reads, clone sequences and assembly contigs with bwa-mem. arXiv preprint arXiv:1303.3997, 2013.

[24] Yanlei Liu, Kai Chen, Lihu Wang, Xinqiang Yu, Chao Xu, Zhili Suo, Shiliang Zhou, Shuo Shi, and Wenpan Dong. Assembly-free reads accurate identification (AFRAID) approach outperforms other methods of DNA barcoding in the walnut family (Juglandaceae). Plant Diversity, 47(1):115–126, January 2025. ISSN 24682659. doi: 10.1016/j.pld.2024.10.002. URL https://linkinghub.elsevier.com/retrieve/pii/S2468265924001677.

[25] Jeffrey M. Marcus. Our love-hate relationship with DNA barcodes, the Y2K problem, and the search for next generation barcodes. AIMS Genetics, 5(1):1–23, 2018. ISSN 2377-1143. doi: 10.3934/genet.2018.1.1. URL http://www.aimspress.com/article/10.3934/genet.2018.1.1.

[26] Daira Melendez, Ali Osman Berk Sapci, Vineet Bafna, and Siavash Mirarab. SPrUCE: Utilizing ultraconserved elements of DNA for population-level genetic diversity estimation. November 2025.

[27] Siavash Mirarab and Vineet Bafna. Analyses of Nuclear Reads Obtained Using Genome Skimming. In Robert DeSalle, editor, DNA Barcoding, volume 2744, pages 247–265. Springer US, New York, NY, 2024. ISBN 978-1-0716-3580-3 978-1-0716-3581-0. doi: 10.1007/978-1-0716-3581-0_16. URL https://link.springer.com/10.1007/978-1-0716-3581-0_16. Series Title: Methods in Molecular Biology.

[28] Zhi-Qiong Mo, Jie Wang, Michael Möller, Jun-Bo Yang, and Lian-Ming Gao. Phylogenetic Relationships and Next-Generation Barcodes in the Genus Torreya Reveal a High Proportion of Misidentified Cultivated Plants. International Journal of Molecular Sciences, 24(17):13216, August 2023. ISSN 1422-0067. doi: 10.3390/ijms241713216. URL https://www.mdpi.com/1422-0067/24/17/13216.

[29] M Nei and W H Li. Mathematical model for studying genetic variation in terms of restriction endonucleases. Proceedings of the National Academy of Sciences, 76(10):5269–5273, October 1979. ISSN 0027-8424, 1091-6490. doi: 10.1073/pnas.76.10.5269. URL https://pnas.org/doi/full/10.1073/pnas.76.10.5269.

[30] Masatoshi Nei. Analysis of gene diversity in subdivided populations. Proceedings of the national academy of sciences, 70(12):3321–3323, 1973.

[31] Rasmus Nielsen and John Wakeley. Distinguishing Migration From Isolation: A Markov Chain Monte Carlo Approach. Genetics, 158(2):885–896, June 2001. ISSN 1943-2631. doi: 10.1093/genetics/158.2.885. URL https://academic.oup.com/genetics/article/158/2/885/6049614.

[32] Koodali T Nishant, Nadia D Singh, and Eric Alani. Genomic mutation rates: what high-throughput methods can tell us. Bioessays, 31(9):912–920, 2009.

[33] Jari Oksanen, Gavin L. Simpson, F. Guillaume Blanchet, Roeland Kindt, Pierre Legendre, Peter R. Minchin, R.B. O’Hara, Peter Solymos, M. Henry H. Stevens, Eduard Szoecs, Helene Wagner, Matt Barbour, Michael Bedward, Ben Bolker, Daniel Borcard, Tuomas Borman, Gustavo Carvalho, Michael Chirico, Miquel De Caceres, Sebastien Durand, Heloisa Beatriz Antoniazi Evangelista, Rich FitzJohn, Michael Friendly, Brendan Furneaux, Geoffrey Hannigan, Mark O. Hill, Leo Lahti, Cameron Martino, Dan McGlinn, Marie-Helene Ouellette, Eduardo Ribeiro Cunha, Tyler Smith, Adrian Stier, Cajo J.F. Ter Braak, and James Weedon. vegan: Community Ecology Package, 2026. URL https://vegandevs.github.io/vegan/. R package version 2. 8–0.

[34] Brian D Ondov, Todd J Treangen, Páll Melsted, Adam B Mallonee, Nicholas H Bergman, Sergey Koren, and Adam M Phillippy. Mash: fast genome and metagenome distance estimation using MinHash. Genome Biology, 17(1):132, December 2016. ISSN 1474-760X. doi: 10.1186/s13059-016-0997-x.

[35] Henrique Miguel Pereira, Simon Ferrier, Michele Walters, Gary N Geller, Rob HG Jongman, Robert J Scholes, Michael William Bruford, Neil Brummitt, Stuart HM Butchart, AC Cardoso, et al. Essential biodiversity variables. Science, 339(6117):277–278, 2013.

[36] N. Tessa Pierce, Luiz Irber, Taylor Reiter, Phillip Brooks, and C. Titus Brown. Large-scale sequence comparisons with sourmash. F1000Research, 8:1006, July 2019. ISSN 2046-1402. doi: 10.12688/f1000research.19675.1. URL https://f1000research.com/articles/8-1006/v1.

[37] Donald L. J. Quicke, M. Alex Smith, Daniel H. Janzen, Winnie Hallwachs, Jose Fernandez-Triana, Nina M. Laurenne, Alejandro Zaldívar-Riverón, Mark R. Shaw, Gavin R. Broad, Seraina Klopfstein, Scott R. Shaw, Jan Hrcek, Paul D. N. Hebert, Scott E. Miller, Josephine J. Rodriguez, James B. Whitfield, Michael J. Sharkey, Barbara J. Sharanowski, Reijo Jussila, Ian D. Gauld, Douglas Chesters, and Alfried P. Vogler. Utility of the DNA barcoding gene fragment for parasitic wasp phylogeny (Hymenoptera: Ichneumonoidea): data release and new measure of taxonomic congruence. Molecular Ecology Resources, 12(4):676–685, July 2012. ISSN 1755-098X, 1755-0998. doi: 10.1111/j.1755-0998.2012.03143.x. URL https://onlinelibrary.wiley.com/doi/10.1111/j.1755-0998.2012.03143.x.

[38] Eleonora Rachtman, Metin Balaban, Vineet Bafna, and Siavash Mirarab. The impact of contaminants on the accuracy of genome skimming and the effectiveness of exclusion read filters. Molecular Ecology Resources, 20(3):1755–0998.13135, May 2020. ISSN 1755-098X. doi: 10.1111/1755-0998.13135. URL https://onlinelibrary.wiley.com/doi/abs/10.1111/1755-0998.13135.

[39] Eleonora Rachtman, Shahab Sarmashghi, Vineet Bafna, and Siavash Mirarab. Quantifying the uncertainty of assembly-free genome-wide distance estimates and phylogenetic relationships using subsampling. Cell Systems, 13(10):817–829.e3, October 2022. ISSN 24054712. doi: 10.1016/j.cels.2022.06.007. URL https://linkinghub.elsevier.com/retrieve/pii/S2405471222002770.

[40] Mahmudur Rahman Hera and David Koslicki. Estimating similarity and distance using FracMinHash. Algorithms for Molecular Biology, 20(1):8, May 2025. ISSN 1748-7188. doi: 10.1186/s13015-025-00276-8. URL https://almob.biomedcentral.com/articles/10.1186/s13015-025-00276-8.

[41] David H. Reed and Richard Frankham. Correlation between Fitness and Genetic Diversity. Conservation Biology, 17(1):230–237, February 2003. ISSN 0888-8892, 1523-1739. doi: 10.1046/j.1523-1739.2003.01236.x. URL https://conbio.onlinelibrary.wiley.com/doi/10.1046/j.1523-1739.2003.01236.x.

[42] John Reynolds, Bruce S Weir, and C Clark Cockerham. Estimation of the coancestry coefficient: basis for a short-term genetic distance. Genetics, 105(3):767–779, 1983.

[43] Shahab Sarmashghi, Kristine Bohmann, M. Thomas P. Gilbert, Vineet Bafna, and Siavash Mirarab. Skmer: assembly-free and alignment-free sample identification using genome skims. Genome Biology, 20 (1):34, December 2019. ISSN 1474-760X. doi: 10.1186/s13059-019-1632-4. URL https://genomebiology.biomedcentral.com/articles/10.1186/s13059-019-1632-4.

[44] Shahab Sarmashghi, Metin Balaban, Eleonora Rachtman, Behrouz Touri, Siavash Mirarab, and Vineet Bafna. Estimating repeat spectra and genome length from low-coverage genome skims with RESPECT. PLOS Computational Biology, 17(11):e1009449, November 2021. ISSN 1553-7358. doi: 10.1371/journal.pcbi.1009449. URL https://dx.plos.org/10.1371/journal.pcbi.1009449.

[45] Jim Shaw and Yun William Yu. Fast and robust metagenomic sequence comparison through sparse chaining with skani. Nature Methods, September 2023. ISSN 1548-7091, 1548-7105. doi: 10.1038/s41592-023-02018-3. URL https://www.nature.com/articles/s41592-023-02018-3.

[46] Montgomery Slatkin. Isolation by distance in equilibrium and non-equilibrium populations. Evolution, 47(1):264–279, 1993.

[47] Kujin Tang, Jie Ren, and Fengzhu Sun. Afann: bias adjustment for alignment-free sequence comparison based on sequencing data using neural network regression. Genome Biology, 20(1):266, December 2019. ISSN 1474-760X. doi: 10.1186/s13059-019-1872-3. URL https://genomebiology.biomedcentral.com/articles/10.1186/s13059-019-1872-3.

[48] Bruce S Weir and C Clark Cockerham. Estimating f-statistics for the analysis of population structure. evolution, pages 1358–1370, 1984.

[49] Hilde M. Wilkinson-Herbots. Genealogy and subpopulation differentiation under various models of population structure. Journal of Mathematical Biology, 37(6):535–585, December 1998. ISSN 0303-6812, 1432-1416. doi: 10.1007/s002850050140. URL http://link.springer.com/10.1007/s002850050140.

[50] Derrick E. Wood, Jennifer Lu, and Ben Langmead. Improved metagenomic analysis with Kraken 2. Genome Biology, 20(1):257, December 2019. ISSN 1474-760X. doi: 10.1186/s13059-019-1891-0. URL https://genomebiology.biomedcentral.com/articles/10.1186/s13059-019-1891-0.

[51] Shengyong Xu, Shijun Xiao, Shilin Zhu, Xiaofei Zeng, Jing Luo, Jiaqi Liu, Tianxiang Gao, and Nansheng Chen. A draft genome assembly of the chinese sillago (sillago sinica), the first reference genome for sillaginidae fishes. GigaScience, 7(9):giy108, 2018.

[52] Le Zhang, Yi-Wei Huang, Jia-Lin Huang, Ji-Dong Ya, Meng-Qing Zhe, Chun-Xia Zeng, Zhi-Rong Zhang, Shi-Bao Zhang, De-Zhu Li, Hong-Tao Li, and Jun-Bo Yang. DNA barcoding of Cymbidium by genome skimming: Call for next-generation nuclear barcodes. Molecular Ecology Resources, 23(2): 424–439, February 2023. ISSN 1755-098X, 1755-0998. doi: 10.1111/1755-0998.13719. URL https://onlinelibrary.wiley.com/doi/10.1111/1755-0998.13719.

[53] Xiang Zhao, Tianlun Zheng, Tianxiang Gao, and Na Song. Whole-genome resequencing reveals genetic diversity and selection signals in warm temperate and subtropical sillago sinica populations. BMC genomics, 24(1):547, 2023.

[54] Ali Osman Berk$apci, Eleonora Rachtman, and Siavash Mirarab. CONSULT-II: Accurate taxonomic identification and profiling using locality-sensitive hashing. Bioinformatics, page btae150, March 2024. ISSN 1367-4811. doi: 10.1093/bioinformatics/btae150. URL https://academic.oup.com/bioinformatics/advance-article/doi/10.1093/bioinformatics/btae150/7630488.

