## Supplemental File 1 for "DipSkmer: Reference-free population genomics with diploid genome skims"

### SA Supplementary mathematical derivations

#### SA.1 Computing Coverage

Recall:

- $L :=$ Length of genome (counting each copy of a chromosome only once),
- $N :=$ number of base pairs in a sample genome skim,
- $\ell :=$ read length,
- $\lambda := \frac{N}{2L}\frac{\ell-k+1}{\ell}$ the $k$-mer coverage.
- each $k$-mer has probability $\rho := (1-\epsilon)^k \simeq e^{-k\epsilon}$ of being error-free.
- $\xi := \lambda\rho$ is the error-free coverage of $k$-mers.

Our calculations will use the repeat spectrum of each sample. Below, we focus on only one sample, omitting the subscript that signifies each sample.

- $M_i :=$ the number of $k$-mers observed exactly $i$ times in a sample.
- $\mathcal{M} :=$ the set of all unique kmers in a diploid genome of length $L$
- $M := |\mathcal{M}|$. note $M \geq L$.
- $\mathcal{M}_0 :=$ the set of homozygote $k$-mers in the genome
  $\mathcal{M}_1 :=$ the set of heterozygote $k$-mers in the genome.
- $M_0 := |\mathcal{M}_0|$ and $M_1 := |\mathcal{M}_1|$. Note: $M = M_0 + M_1$ and $L = M_0 + \frac{1}{2}M_1$.

For a genome skim

$$
\begin{aligned}
\mathbb{E}[M_0] &= L(1-\theta)^k \\
\mathbb{E}[M_1] &= 2L\Big(1-(1-\theta)^k\Big) \\
\mathbb{E}[M] &= L(1-\theta)^k + 2L\Big(1-(1-\theta)^k\Big) = L(2-(1-\theta)^k)
\end{aligned}
$$

Note that $M$ increases with increasing $\theta$. To sample a k-mer at random, we first select a location at random, and next, we pick a k-mer from one of the two haplotypes. Thus,

$$
\begin{aligned}
\Pr(x \in \mathcal{M}_0 | x \in \mathcal{M}) &= \frac{(1-\theta)^k}{(1-\theta)^k + 2(1-(1-\theta)^k)} = \frac{(1-\theta)^k}{2-(1-\theta)^k} \\
\Pr(x \in \mathcal{M}_1 | x \in \mathcal{M}) &= \frac{2-2(1-\theta)^k}{2-(1-\theta)^k}\ .
\end{aligned}
$$

Recall that $\rho = (1-\epsilon)^k \approx \epsilon^{-k}$ and $\xi = \lambda\rho$. Let

$$A_{x,i} = \text{ Event that a genomic kmer } x \text{ appears } i \text{ times error-free in a sample.}$$

Then,

$$\begin{aligned}\Pr(A_{x,i}|x \in \mathcal{M}) &= \underbrace{\Pr(A_{x,i}|x \in \mathcal{M}_0)}_{\text{Poisson with rate } 2\xi} \Pr(x \in \mathcal{M}_0|x \in \mathcal{M}) + \underbrace{\Pr(A_{x,i}|x \in \mathcal{M}_1)}_{\text{Poisson with rate } \xi} \Pr(x \in \mathcal{M}_1|x \in \mathcal{M}) \\ &= \frac{e^{-2\xi}\xi^i \left(\left(2^i - 2e^{\xi}\right)(1-\theta)^k + 2e^{\xi}\right)}{i!\left(2-(1-\theta)^k\right)}.\end{aligned} \tag{S1}$$

A family of equations can be obtained by taking the ratio of consecutive counts of error-free k-mers as

$$\nu_i = \frac{\Pr(A_{x,i+1}|x \in \mathcal{M})}{\Pr(A_{x,i}|x \in \mathcal{M})} = \frac{2\xi}{(i+1)} \left( \frac{e^{\xi} - \left(e^{\xi} - 2^i\right)(1-\theta)^k}{2e^{\xi} + \left(2^i - 2e^{\xi}\right)(1-\theta)^k} \right) \tag{S2}$$

for $i \geq 2$. We aim to estimate two unknowns, $\xi$ and $\theta$, which is possible if we use any two equations from this family. In practice, using two consecutive equations leads to a simplification. Note that

$$2\frac{(i+2)\nu_{i+1}}{2\xi} = 3 - \frac{2\xi}{\nu_i(i+1)} \implies \nu_i = -\frac{2\xi^2}{(i+1)(i+2)\nu_{i+1} - 3(i+1)\xi}$$

which completely eliminates $\theta$, and allows us to arrive at an estimate of $\xi$:

$$\hat{\xi} = \frac{3}{4}(i+1)\nu_i - \frac{1}{4}\sqrt{i+1}\sqrt{\nu_i}\sqrt{9(i+1)\nu_i - 8(i+2)\nu_{i+1}} \tag{S3}$$

With $\hat{\xi}$ estimated, we then use Eq. (S2) to estimate $\theta$ directly. It can be confirmed that

$$\hat{\theta} = 1 - \left(\frac{2e^{\xi}(\xi - \nu_i(1+i))}{\nu_i(1+i)(2^i - 2e^{\xi}) + 2\xi(e^{\xi} - 2^i)}\right)^{1/k}$$

satisfies (S2), giving us an estimator for $\theta$. While any $i$ can be used, we follow Skmer[43] and set it to $\arg\max_{i>1} \hat{M}_i$ (i.e., the first mode of the skim's $k$-mer repeat spectrum).

Note that when $9(i+1)\nu_i < 8(i+2)\nu_{i+1}$, we have no solutions , and a different strategy must be employed. In these situations, we simply assume a fixed $\theta^*$ (set to 0.005 by default), which is set *a priori* and is not estimated from the data. Then, we can use (S2) directly to estimate $\xi$ by numerical optimization:

$$\hat{\xi} = \arg\min_{\xi} \left(\nu_i - \frac{2\xi}{1+i}\left(\frac{e^{\xi} - (e^{\xi} - 2^i)(1-\theta^*)^k}{2e^{\xi} + (2^i - 2e^{\xi})(1-\theta^*)^k}\right)\right)^2$$

Regardless of which method is used to compute $\hat{\xi}$, we next need to decompose it into its two components $\lambda$ and $\rho = (1-\epsilon)^k$. To do so, we cannot use the family of equations given by (S2) because they are only a function of $\xi$. Instead, we need to examine error-prone $k$-mers, and in particular, $i = 1$, where the error is concentrated. It can be checked that:

$$\sum_{i=2}^{\infty} i \Pr(A_{x,i}|x \in \mathcal{M}) = \frac{2e^{-2\xi}\left(e^{\xi}-1\right)\xi\left((1-\theta)^k + e^{\xi}\right)}{2-(1-\theta)^k}$$

Let $\hat{M}_i =$ the count of $k$-mers seen $i$ times in the presence of error. Note that

$$\mathbb{E}[\sum_{i=1}^{\infty} i\hat{M}_i] = 2L\lambda = 2\lambda \frac{\mathbb{E}[M]}{2-(1-\theta)^k}$$

Then, assuming that all erroneous $k$-mers fall under $\hat{M}_1$,

$$\begin{aligned}\mathbb{E}[\hat{M}_i] &= \begin{cases} \mathbb{E}[\sum_{x\in\mathcal{M}} \Pr(A_{x,i}|x \in \mathcal{M})] & i > 1 \\ \mathbb{E}[\sum_{i=1}^{\infty} i\hat{M}_i - \sum_{i=2}^{\infty} i\hat{M}_i] & i = 1 \end{cases} = \begin{cases} \mathbb{E}[\sum_{x\in\mathcal{M}} \Pr(A_{x,i}|x \in \mathcal{M})] & i > 1 \\ 2\lambda\frac{\mathbb{E}[M]}{2-(1-\theta)^k} - \mathbb{E}[\sum_{i=2}^{\infty} i\hat{M}_i] & i = 1 \end{cases} \\ &= \begin{cases} \mathbb{E}[M]\frac{e^{-2\xi}\xi^i\left(\left(2^i-2e^{\xi}\right)(1-\theta)^k+2e^{\xi}\right)}{i!(2-(1-\theta)^k)} & i \geq 2 \\ \mathbb{E}[M]\frac{2\left(\lambda-e^{-2\xi}\left(e^{\xi}-1\right)\xi\left((1-\theta)^k+e^{\xi}\right)\right)}{2-(1-\theta)^k} & i = 1 \end{cases}\end{aligned} \tag{S4}$$

For dividing $\xi$ to $\epsilon$ and $\lambda$, we can use

$$\frac{\hat{M}_1}{\hat{M}_i} = 2i!\xi^{-i}\frac{\xi\left(1-e^{\xi}\right)(1-\theta)^k + e^{\xi}\left(\xi - e^{\xi}(\xi-\lambda)\right)}{\left(2^i - 2e^{\xi}\right)(1-\theta)^k + 2e^{\xi}}$$

which can be solved to get:

$$\hat{\lambda} = \frac{\hat{M}_1}{\hat{M}_i}\frac{\hat{\xi}^i\left(\left(2^i e^{-2\hat{\xi}} - 2e^{-\hat{\xi}}\right)(1-\theta)^k + 2e^{-\hat{\xi}}\right)}{2i!} + \hat{\xi}\left(e^{\hat{\xi}}-1\right)\left(e^{-2\hat{\xi}}(1-\theta)^k + e^{-\hat{\xi}}\right), \quad \hat{\epsilon} = 1 - (\hat{\xi}/\hat{\lambda})^{1/k}\ .$$

#### SA.2 Extra nodes on diploid estimate of $\theta$

##### SA.2.1 Limitations of Intersection

From the table in Section 4.3, we can compute:

$$\mathbb{E}[I] = \mathbf{P}\cdot\mathbf{I} = \frac{1}{11}\Big(\eta_2(2)\big(11Q\eta_1(2) - 4(Q-1)\eta_1(1)\big) - 4(Q-1)\eta_2(1)\big(\eta_1(1)+\eta_1(2)\big)\Big) \tag{S5}$$

$$\mathbb{E}[U] = \mathbf{P}\cdot\mathbf{U} = \frac{(1-Q)}{11}\Big(-4(\eta_2(1)\eta_1(1)+\eta_2(1)\eta_1(2)+\eta_1(1)\eta_2(2))+12(\eta_2(1)+\eta_1(1))+5(\eta_2(2)+\eta_1(2))\Big)+ Q\Big(\eta_2(2)+\eta_1(2)-\eta_2(2)\eta_1(2)\Big) \tag{S6}$$

Recall that for the case of $t = 1$, we can replace $\eta_1(1)$ with $\eta_1 = (1 - e^{\xi_1})$, and note that:

$$\eta_1(2) = (1-e^{2\xi_1}) = (1-e^{\xi_1})(1+e^{\xi_1}) = (1-e^{\xi_1})(2-(1-e^{\xi_1})) = \eta_1(2-\eta_1) \tag{S7}$$

(ditto for $\eta_2(2)$). Then, we can derive:

$$Q = \frac{11\mathbb{E}[I] + 4\eta_2\eta_1(\eta_1+\eta_2-5)}{\eta_1\eta_2\left(11\eta_2\eta_1 - 18(\eta_1+\eta_2)+24\right)}\ .$$

which can be compared with the equation using Jaccard after replacing $\eta_1(1)$ with $\eta_1 = 1 - e^{-\xi_1}$ and $\eta_1(2) = \eta_1(2-\eta_1)$ and further simplifications (for $t = 1$):

$$\mathbb{E}[I] = \frac{6}{11}\eta_1\eta_2\left(\frac{11Q}{6}\eta_1\eta_2 - \left(3Q+\frac{2}{3}\right)(\eta_1+\eta_2)+4Q+\frac{10}{3}\right)$$

$$Q = \frac{4\eta_1\eta_2(\eta_1+\eta_2-5) + J\left(2(2\eta_1\eta_2+11)(\eta_2+\eta_1) - 5(\eta_1+\eta_2)^2 - 10\eta_1\eta_2 + 11(\psi_1+\psi_2)\right)}{(J+1)\eta_2\eta_1\left(11\eta_2\eta_1 - 18\eta_2 - 18\eta_1 + 24\right) + 6J(\eta_2^2+\eta_1^2)} \tag{S8}$$

The equation using intersection alone is much cleaner, and it is tempting to use intersection instead of Jaccard. However, a closer examination reveals that using intersection alone, as opposed to Jaccard, which depends on both intersection and union, is not enough to compute distance. In fact, a surprising effect can be proved with ease.

**Remark S1** *Intersection size $I$ is not enough to compute distances uniquely. For $\eta_1 = \eta_2 = \frac{2}{11}\left(9-\sqrt{15}\right)$ (roughly, 3.6× per genome with 0.003 error), $\mathbb{E}[I]$ is not a function of $Q$.*

More generally, the figure below shows that the intersection can change very slowly with $Q$, making it a highly noisy estimator. Thus, the estimation of distance should use the Jaccard index instead of the Union.

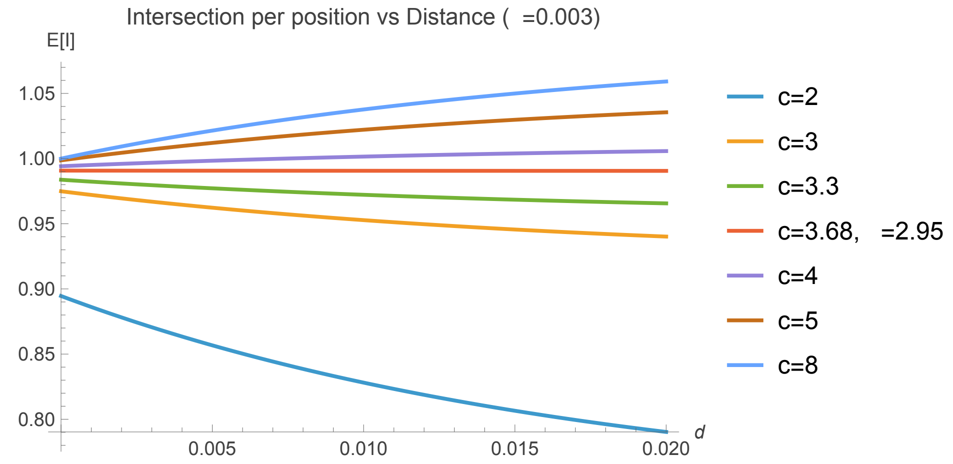


##### SA.2.2 Haploid to Diploid conversion.

We can go from a Haploid Skmer-1 estimate to a diploid estimate by noting:

$$J = \frac{(1-\hat{d})^k}{2-(1-\hat{d})^k}$$

and plugging this $J$ back in Equation (2), we get:

$$\hat{\theta}(\hat{d}) = 1 - \left( 2 \frac{\frac{11(1-\hat{d})^k}{(1-\hat{d})^k-2} + 6}{\frac{11(1-\hat{d})^k}{(1-\hat{d})^k-2} + 1} \right)^{\frac{6}{11}\frac{1}{k}} = 1 - \left( \frac{11}{6} \left( 1 - \frac{6-1}{6(1-\hat{d})^k - 1} \right) + 1 \right)^{\frac{6}{11}\frac{1}{k}} \tag{S9}$$

Note that the Taylor expansion of this function around 0 is

$$\frac{6}{5}d + \frac{1671}{25}d^2 + \frac{511496}{125}d^3 + \dots$$

Drawing this equation (Fig. S1b) shows that for small enough $\theta < 0.002$, a simple multiplication by $6/5$ is sufficient for translating from haploid to diploid estimates.

### SB Supplementary Tables

| Method | 0.5× | 1× | 2× | 4× | 6× | 8× |
|---|---|---|---|---|---|---|
| DipSkmer | **25.1%** | **28.4%** | **22.4%** | **26.0%** | **37.6%** | 40.5% |
| ReSkmer-noref | 62.6% | 36.3% | 33.0% | 30.3% | 38.7% | 43.9% |
| Skmer | 49.3% | 46.8% | 49.2% | 64.6% | 169.9% | 229.0% |
| Mash | 159.1% | 97.5% | 528.3% | 310.7% | 328.4% | 268.7% |

Table S1: **MAE of reference-free $\theta$ estimates in Biological Data.** The mean absolute error (MAE) of reference-free $\theta$ estimates with different methods. Each value represents the average error at a specific coverage across all 12 populations. For error calculation, ANGSD as reference $\theta$.

| species | population | $\theta$ (ANGSD) |
|---|---|---|
| *Certhidea fusca* | f-espanola | 0.0006 |
| *Pinaroloxias inornata* | cocos island | 0.0012 |
| *Certhidea fusca* | cristobal | 0.0019 |
| *Geospiza conirostris* | c-espanola | 0.0024 |
| *Geospiza conirostris* | genovesa | 0.0028 |
| *Ovis aries* | panou | 0.0036 |
| *Ovis aries* | oula | 0.0039 |
| *Apis cerana* | niupeng | 0.0054 |
| *Apis cerana* | zhongshui | 0.0056 |
| *Silago sinica* | dongying | 0.0063 |
| *Silago sinica* | qingdao | 0.0064 |
| *Silago sinica* | wenzhou | 0.0068 |

Table S2: **ANGSD Estimates of $\theta$ per Population:** Species, population name, and ANGSD $\theta$ as estimated by the SPRUCE paper[26]. Populations are ordered by increasing $\theta$.

| **Common Name** | **Species** | **Accession** | **Genome Size** | **UR** |
|---|---|---|---|---|
| Club Moss | Selaginella moellendorffii | GCF_000143415.4 | 208,216,466 | 0.42 |
| Algae | *Chondrus crispus* | GCF_000350225.1 | 103,988,016 | 0.51 |
| Oyster | *Ostrea edulis* | GCF_947568905.1* | 98,251,904 | 0.64 |
| Nematode | *Nemopilema nomurai* | GCA_003864495.1 | 210,407,387 | 0.71 |
| Rotifer | *Brachionus plicatilis* | GCA_010279815.1 | 101,125,688 | 0.85 |
| Leech | *Hirudo medicinalis* | GCA_011800805.1 | 155,214,494 | 0.90 |

Table S3: **Assemblies used in simulations.** Genomes used in these simulations have different levels of repetitiveness measured by uniqueness ratio ($UR = {}^{\text{unique } k\text{-mers}}/_{\text{genome size}}$). Only one scaffold (NC_079166.1) of the *O. edulis* assembly is used to provide a uniqueness ratio in between other genomes.

| Genome Size | Species | DipSkmer | Skmer | Mash | Respect | ReSkmer |
|---|---|---|---|---|---|---|
| 0.24 | *Apis cerana* | 37.87 | 39.58 | 10.60 | 554.56 | 46.78 |
| 0.58 | *Silago sinica* | 170.37 | 171.62 | 36.33 | 771.95 | 305.79 |
| 0.76 | *Tautogolabrus adspersus* | 214.77 | 214.98 | 49.28 | 814.64 | 308.12 |
| 1.18 | *Geopsiza conirostris* | 213.39 | 215.03 | 39.48 | 469.28 | 374.39 |
| 2.73 | *Ovies aries* | 969.28 | 970.77 | 179.97 | 1164.45 | 1440.11 |

Table S4: **Query running time (in seconds) comparison across different genome sizes (Gbp).** Genomes were subsampled to 2× coverage and multi-threaded (10 threads). Mash was not multithreaded, since sketching reads cannot be parallelized. All processes were run on an *AMD EPYC 7742 64-Core Processor*. DipSkmer, ReSkmer, and Skmer were run using the "query" command, while runtime for Mash refers to the "sketch" command.

### SC Supplementary Figures

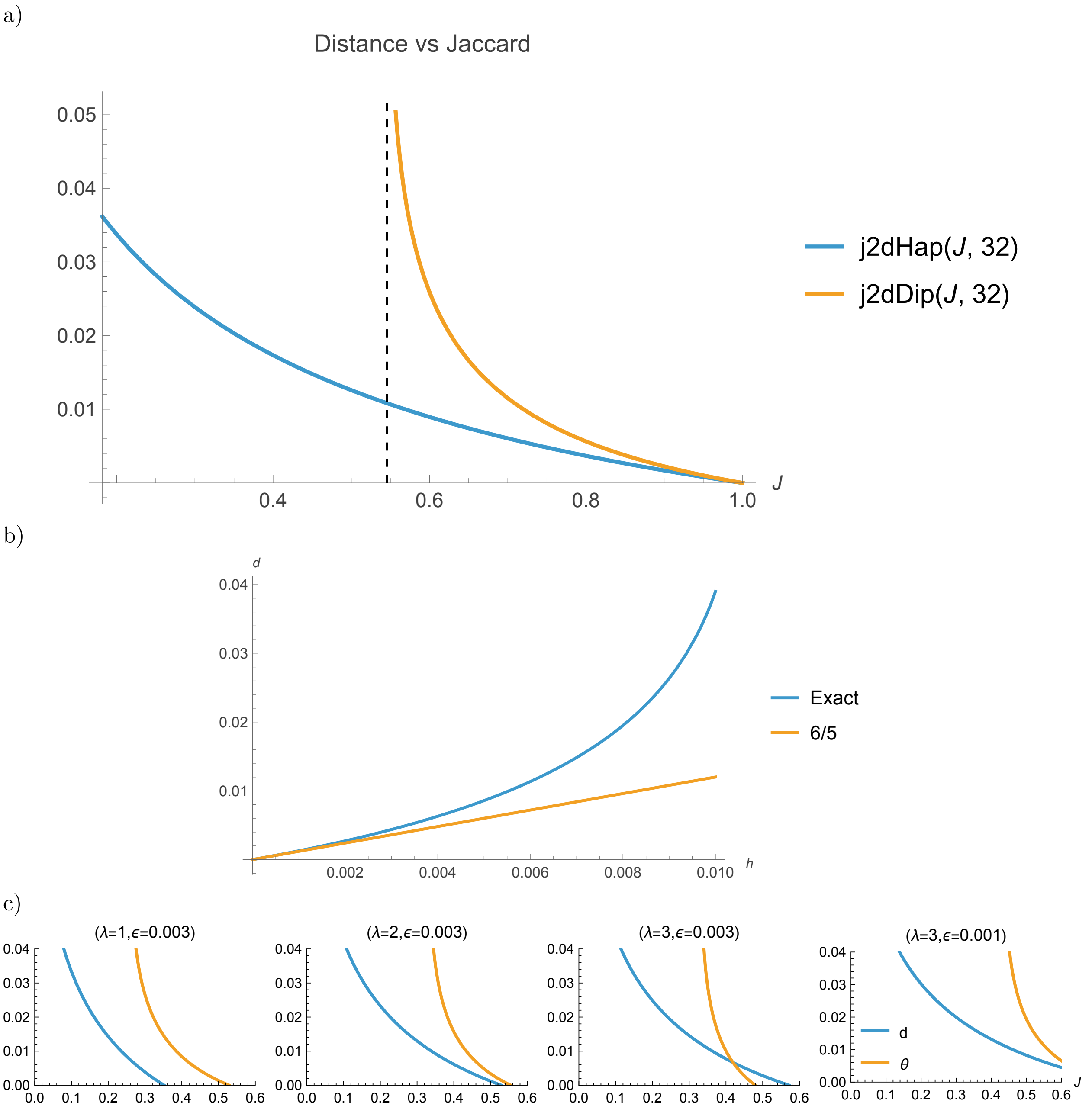


Figure S1: a) The relationship between Jaccard and distance depends on whether we model heterozygosity for diploids or assume a haploid genome. We show $\theta = 1 - \left(\frac{2J}{1+J}\right)^{1/k}$ and $\theta = 1 - \left(\frac{2J-12/11}{1+J-12/11}\right)^{\frac{6}{11}\frac{1}{k}}$ for haploids and diploids, respectively, for $k = 32$. Note that for diploids, the transformation is only possible if $J > \frac{6}{11}$, shown as a dotted line. Recall that the derivation for diploidy relies on independent sites assumptions and thus cannot be used for high distances (e.g., $\theta > 0.02$). b) The shape of Equation (S9) for translating a haploid estimate $h$ from Mash/Skmer (for genomes) to a diploid estimate $d$ from our Eq. (2). Note that for very small $\theta$, simply multiplying by $\frac{6}{5}$ is a good approximation. c) The relationship between Jaccard and $\theta$ changes with different choices of coverage and sequencing error ($d$ for haploid and $\theta$ for diploid).

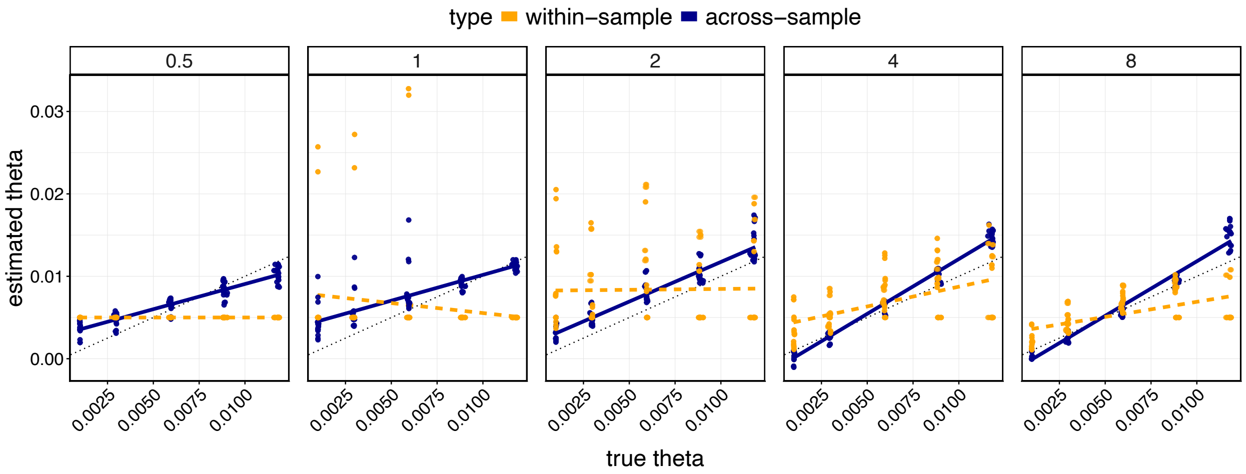


Figure S2: **Within-sample vs. across-sample estimates of $\theta$.** Estimates of $\theta$ ($y$-axis) using Eq. (7) (within-sample) are compared to those obtained across samples using pairs (Section 4.3). Each panel shows a different coverage, and the $x$-axis corresponds to the true $\theta$ value. Note that within-sample estimates completely fail at lower coverages, where a default value of $\theta_* = 0.005$ is used.

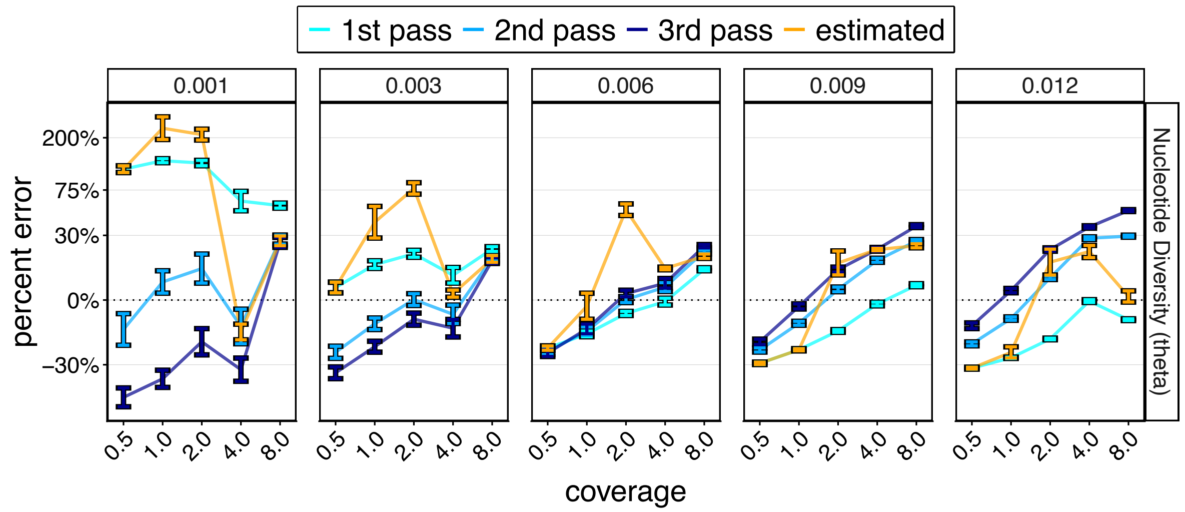


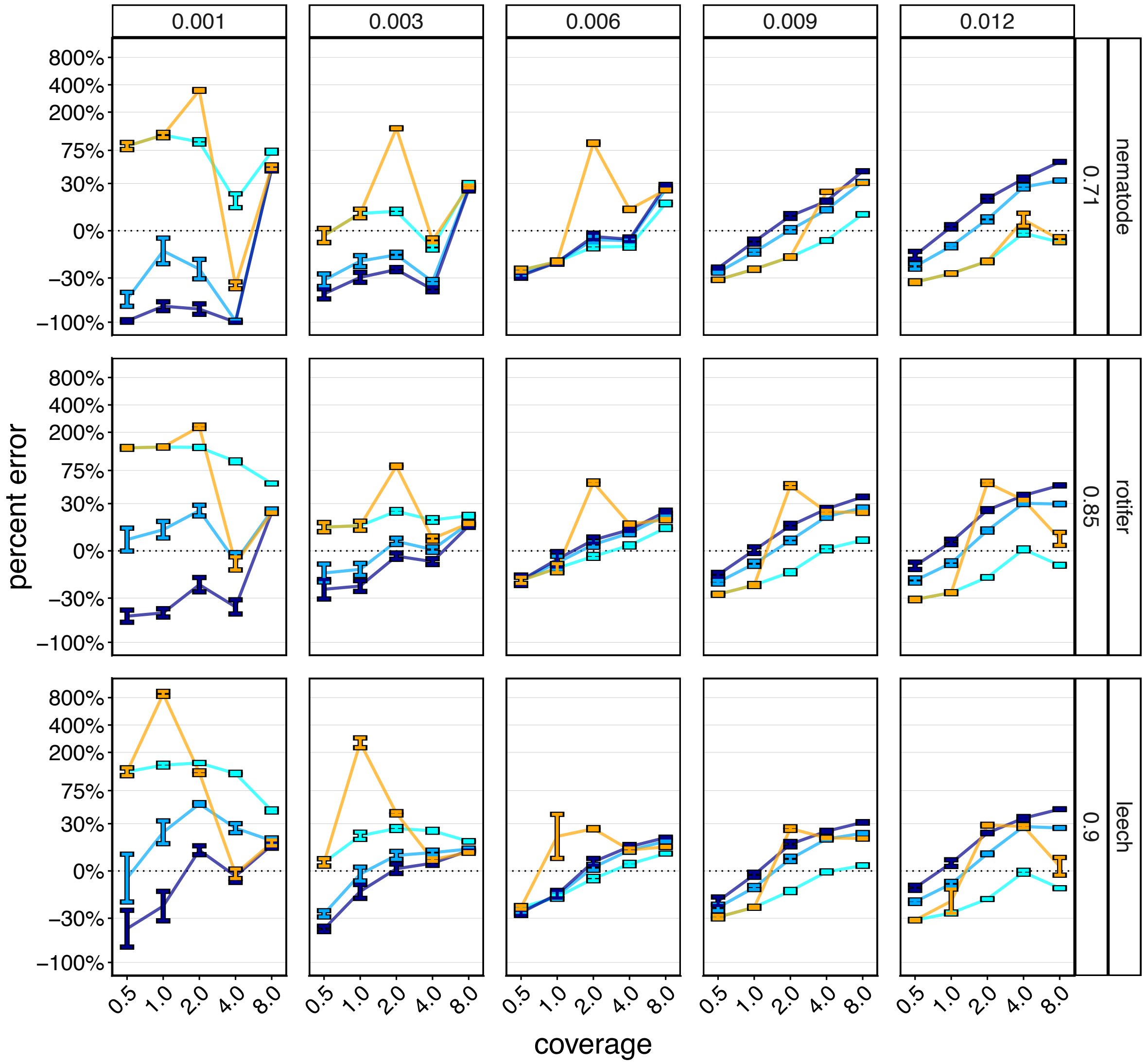

Figure S3: **Iterative vs. ratio-based methods for within-sample $\theta$ estimation in simulated datasets**. Percent error using within-sample estimates using Eq. (7) vs. all passes of the iterative approach. The first panel represents the average of all simulated genomes, while subsequent rows represent individual root genomes with different levels of repetitiveness ($k$-mer uniqueness ratio on the right strip, along with the common name of the genome).

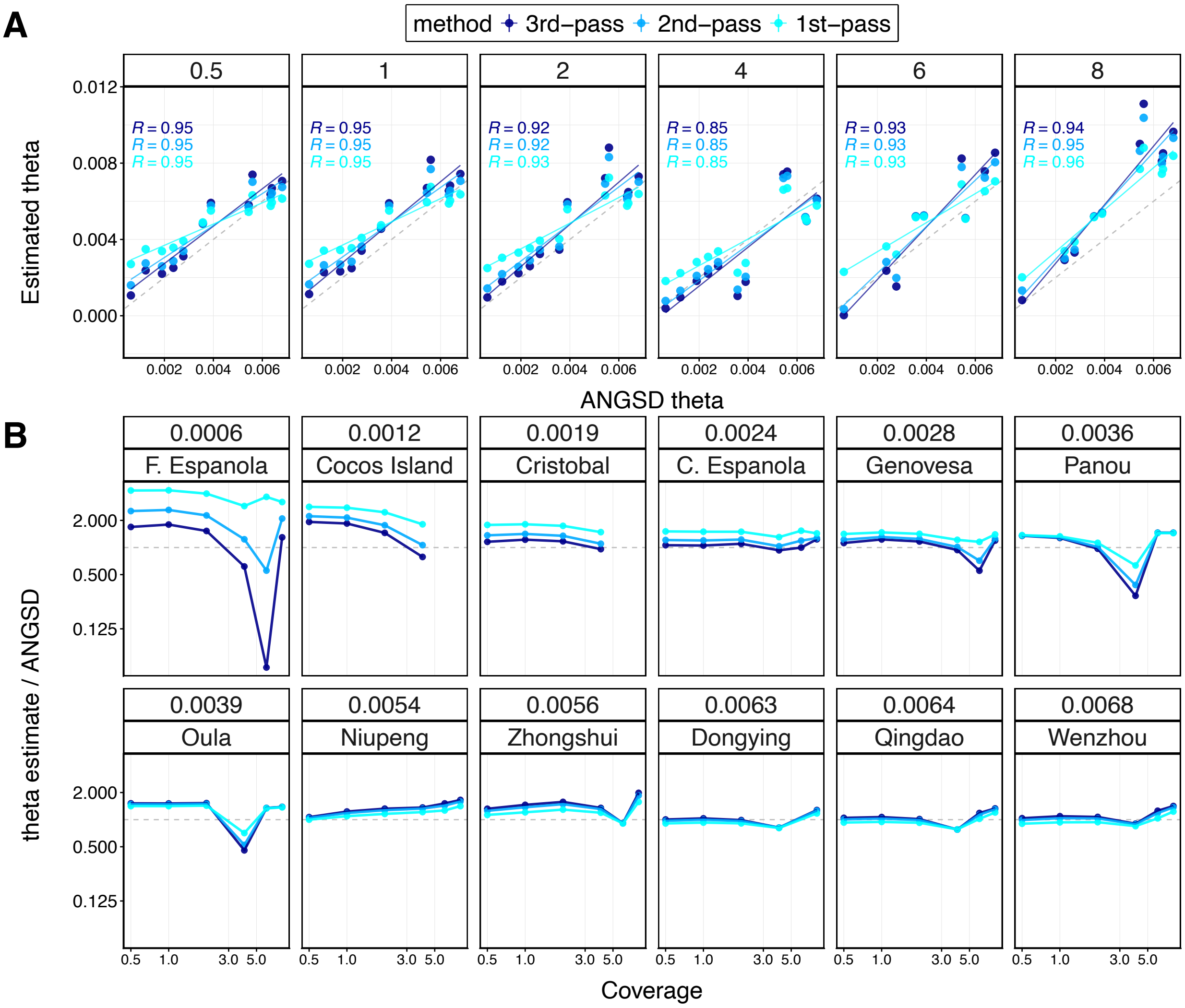


Figure S4: **Results from DipSkmer $\theta$ estimates at different passes per species**. **A)** DipSkmer estimates of $\theta$ ($y$-axis) versus ANGSD $\theta$ estimates ($x$-axis). Darker colors represent later passes, and the dotted line is the unity line. Pearson correlation coefficients are shown. **B)** The ratio of dipskmer $\theta$ to Angsd $\theta$. The dashed line at $y = 1$ represents the ideal case of obtaining an identical estimate.

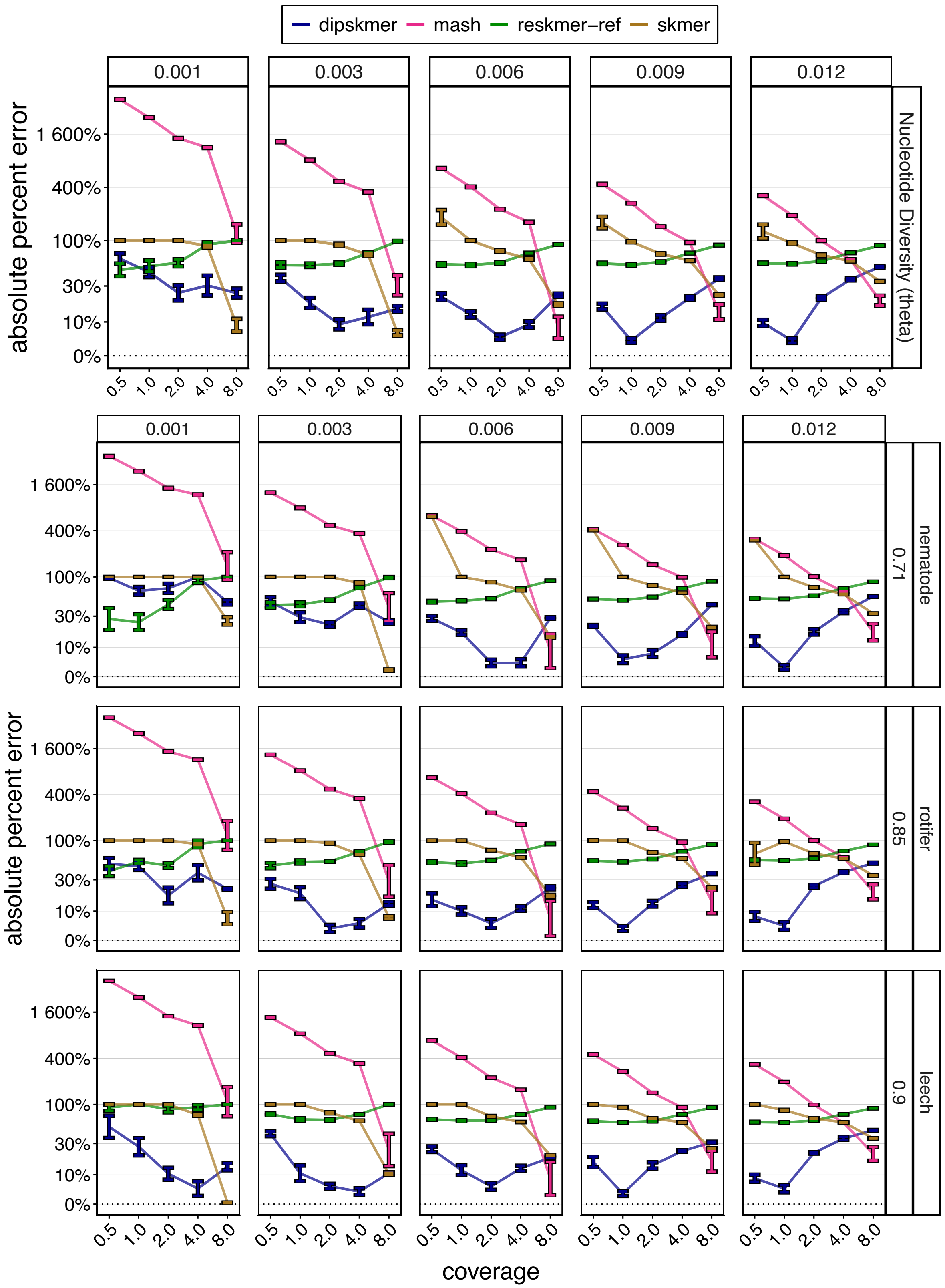


Figure S5: **Absolute Error in Nucleotide Diversity ($\theta$) Estimates. Top Panel:** Range of absolute % error across all simulated genomes and replicates ($y$-axis) plotted against simulated sequencing coverage ($x$-axis). The *Top strip* shows different simulated populations' true $\theta$ values. **Bottom Panel:** Range of absolute % error across all replicates ($y$-axis) for specific genomes with varying levels of $k$-mer repetitiveness (uniqueness ratio and genome name on the *right strip*).

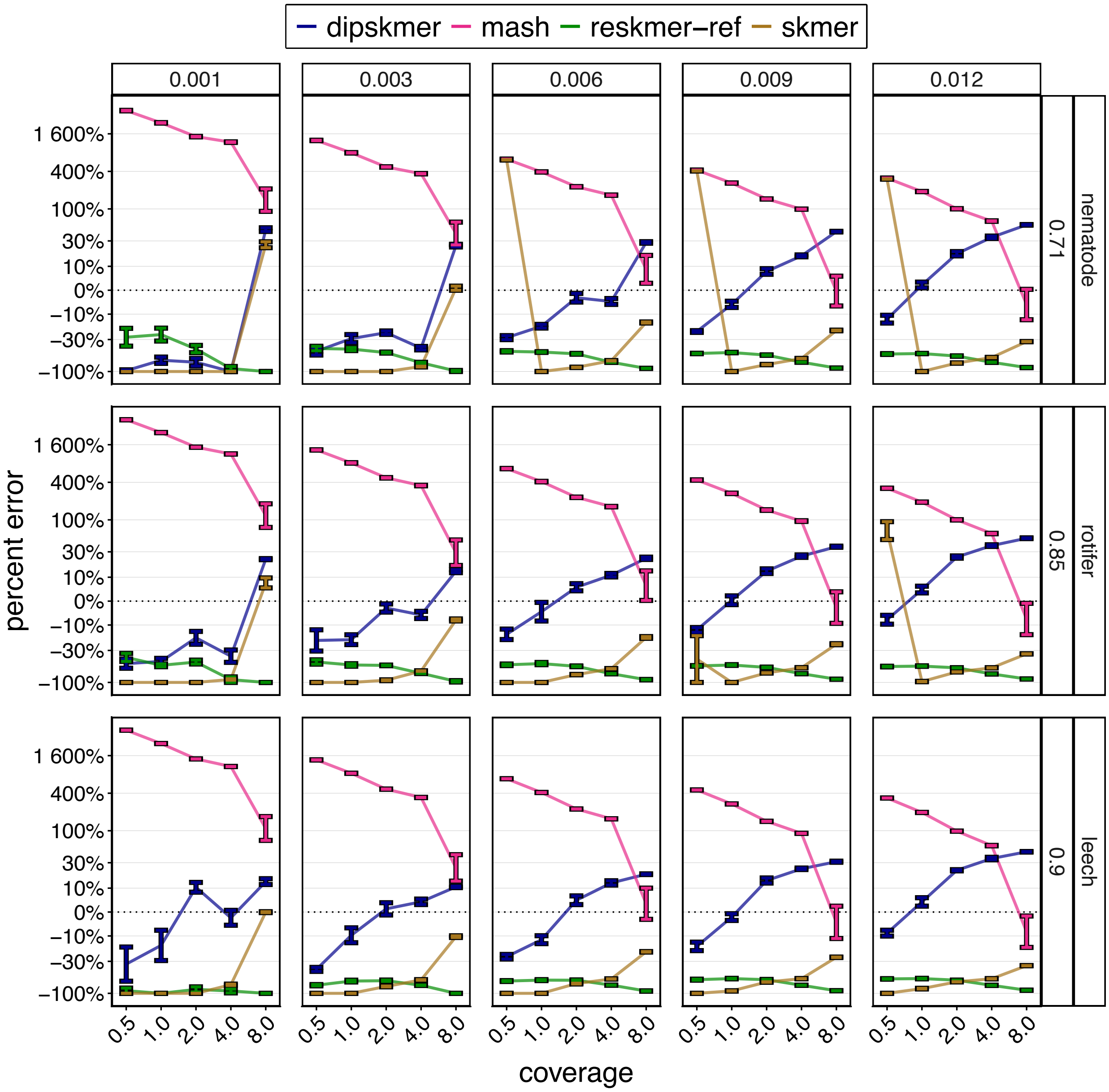


Figure S6: **Effect of Repetitiveness on Reference-Free $\theta$ Estimates.** Percent error ($y$-axis) plotted against coverage ($x$-axis). Columns represent simulated $\theta$ values, and rows show results using genomes with different levels of $k$-mer repetitiveness (uniquess ratio and genome used on the *right strips*).

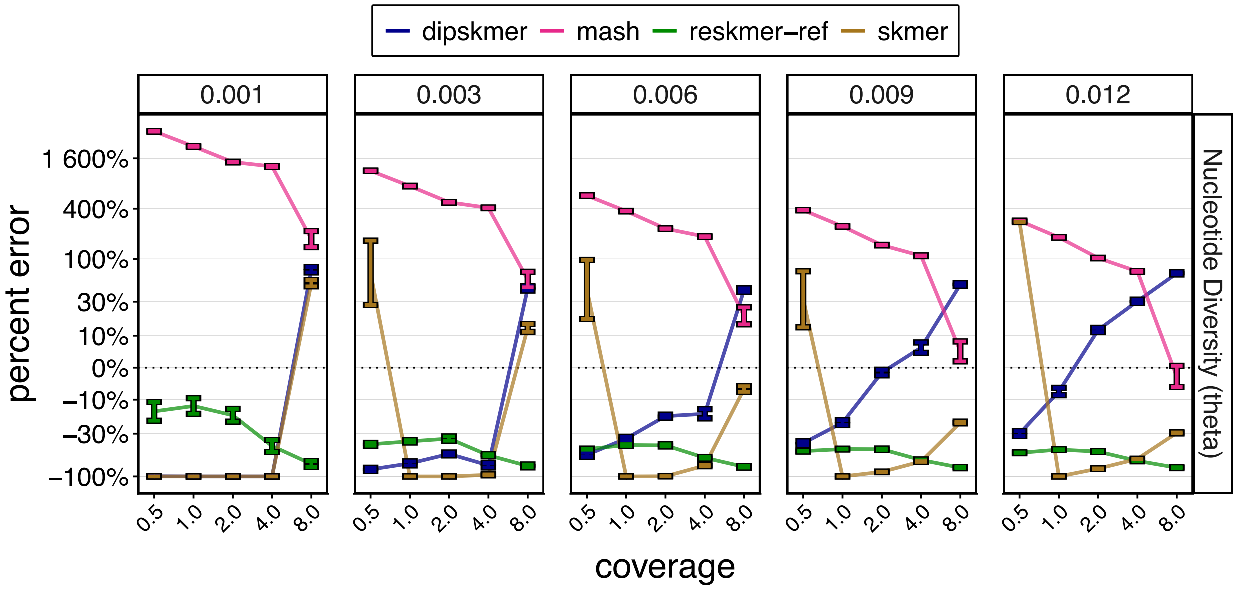


Figure S7: **Percent Error in $\theta$ Estimation with Highly Repetitive Genomes:** We perform the same simulation process described in Methods using three extremely repetitive genomes ($UR \in \{0.6; 0.5; 0.4\}$ – meaning that at least 40% of all $k$-mers appear more than once in the genome). We then estimate $\theta$ using different methods. Error bars represent the range of values from results with all three genomes.

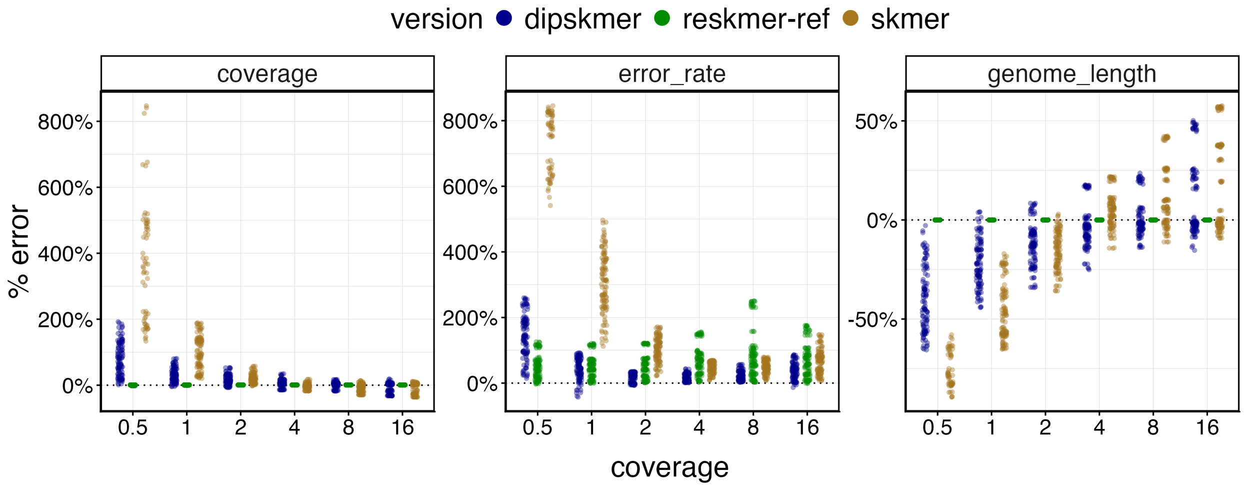


Figure S8: **Sequencing Parameter Estimation Comparison:** Panels represent different sequencing parameter estimates. Percent error ($y$-axis) plotted against simulated coverages ($x$-axis), genome lengths corresponding to those in Table S3, and an error rate of $\sim 0.003\%$ (Phred $= 25$). Note that ReSkmer is given access to the reference genome and therefore has access to the correct genome length and thus coverage.

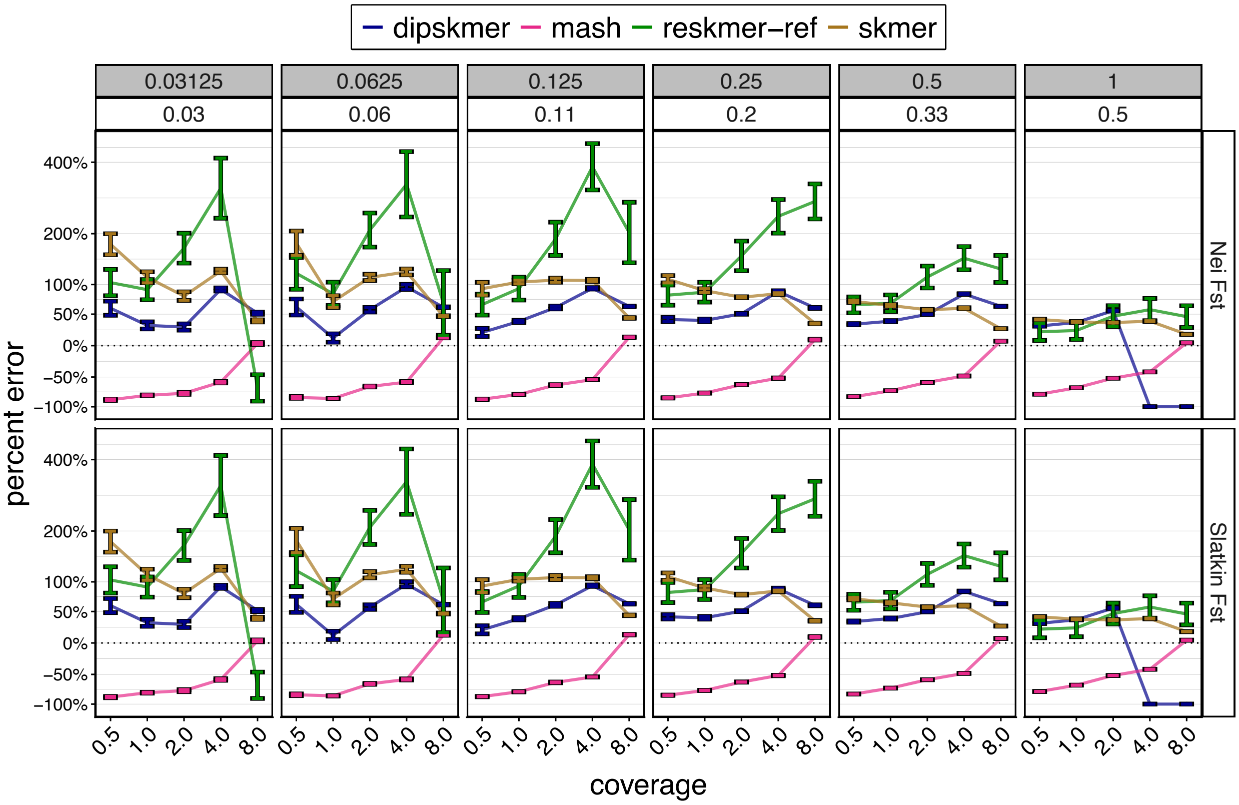


Figure S9: **Percent error across $F_{st}$ definitions.** Rows show different $F_{ST}$ definitions. Slatkin[46] is defined as $1 - \frac{(\theta_x+\theta_y)}{D_{XY}+(\theta_x+\theta_y)}$ , while Nei[30] is defined as $\frac{(\theta_{xy}-(\theta_x+\theta_y)/2)}{\theta_{xy}}$. The top strip shows different divergence time between the simulated populations in coalescent units, under which is the mean true WC $F_{st}$ value across simulations. The $x$-axis represents coverage, while the $y$-axis illustrates the % error of estimates obtained using different methods.

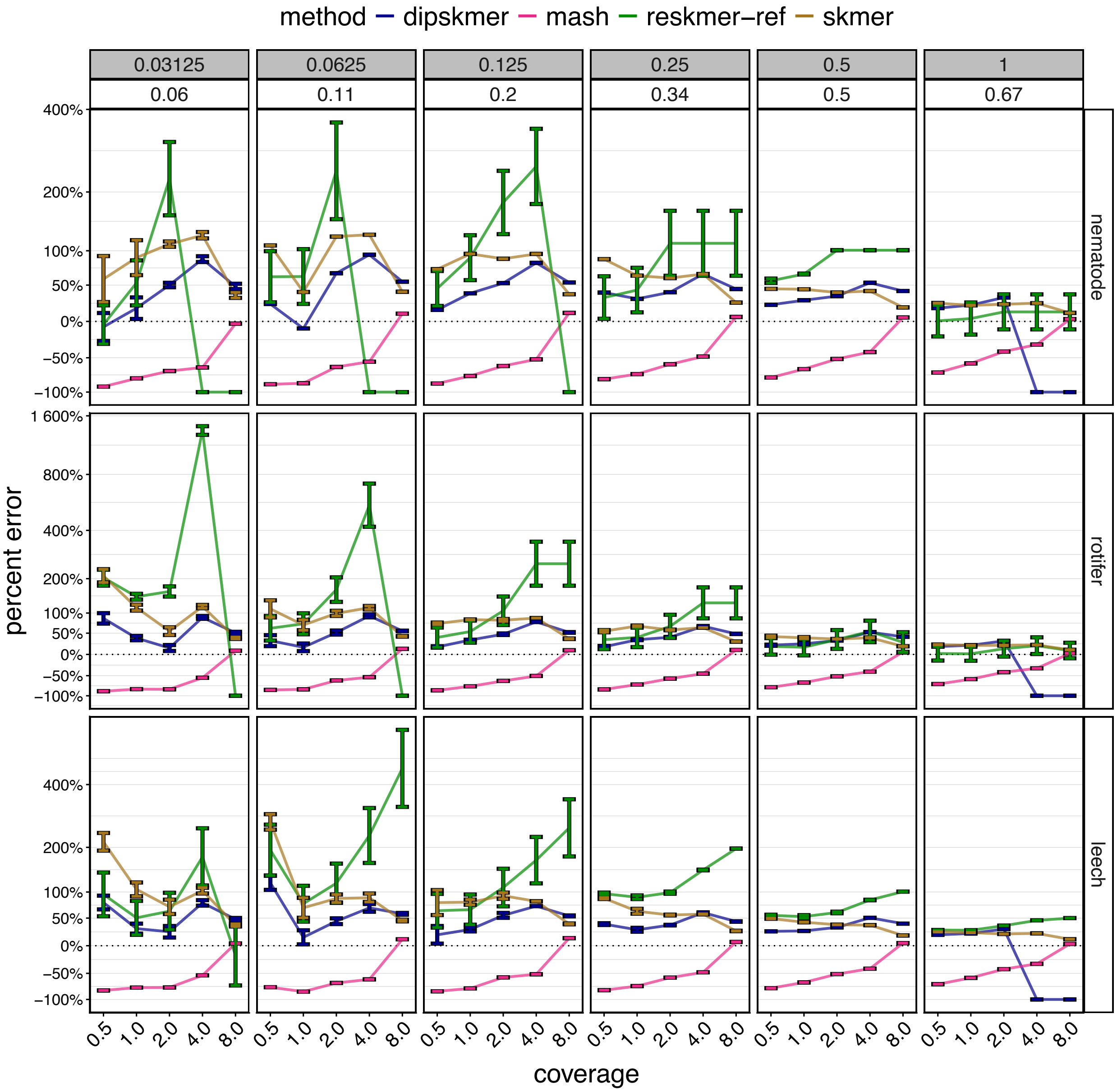


Figure S10: **Weir and Cockerham $F_{st}$ in Variably Repetitive Genomes.** The $x$-axis represents coverage, while the $y$-axis illustrates the % error of Weir and Cockerham $F_{st}$ estimates obtained using different methods. Rows represent different genomes with varying uniqueness ratios (left strip). Columns represent divergence times between simulated populations in coalescent units (gray strip), under which is the mean true WC $F_{st}$.

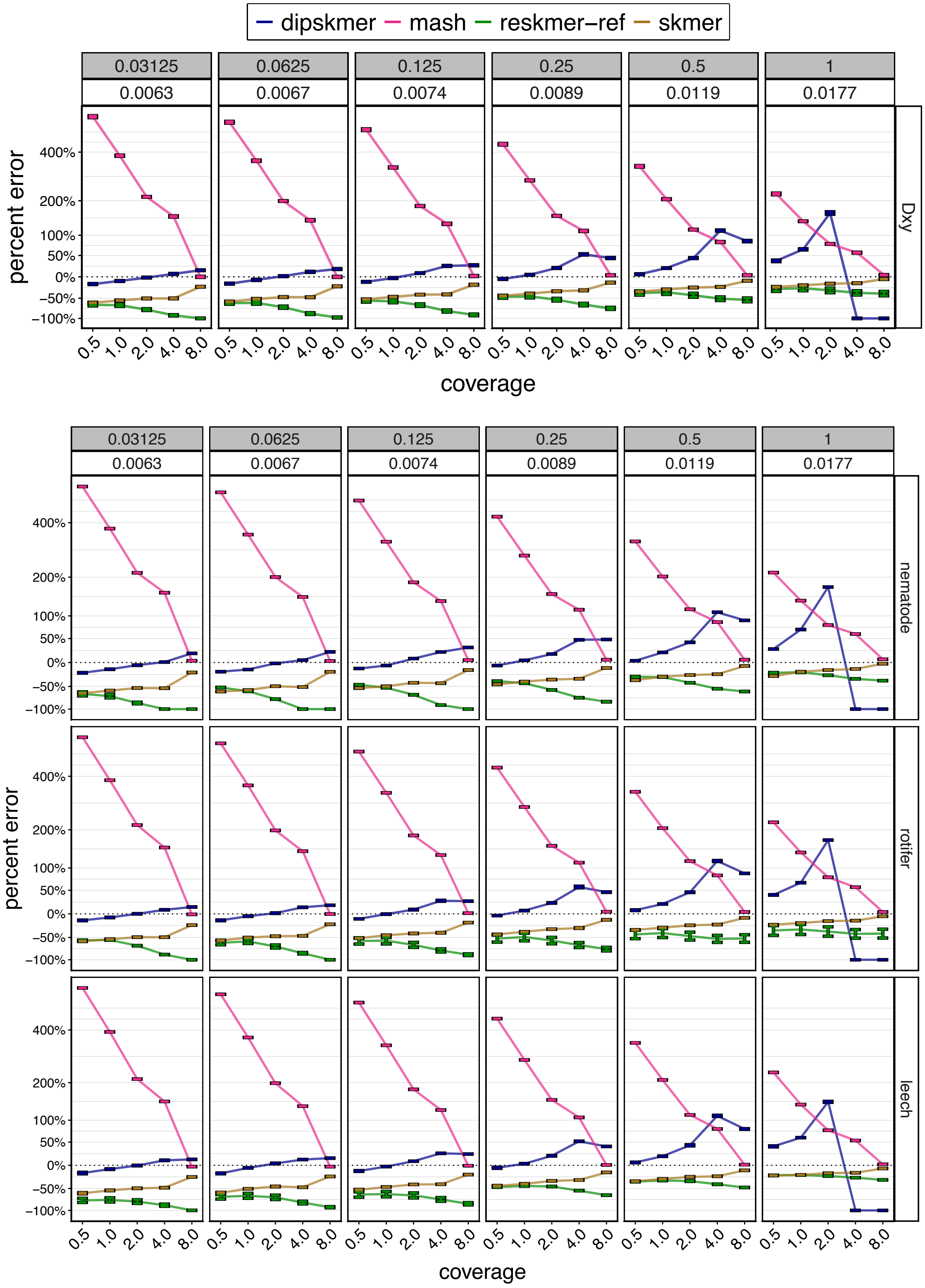


Figure S11: $D_{xy}$ **Estimation Error in Variably Repetitive Genomes**. The $x$-axis represents coverage, while the $y$-axis illustrates the % error of $D_{xy}$ estimates obtained using different methods. Rows represent different genomes with varying uniqueness ratios (left strip). Columns represent divergence times between simulated populations in coalescent units (top strip), under which is the mean true $D_{xy}$.

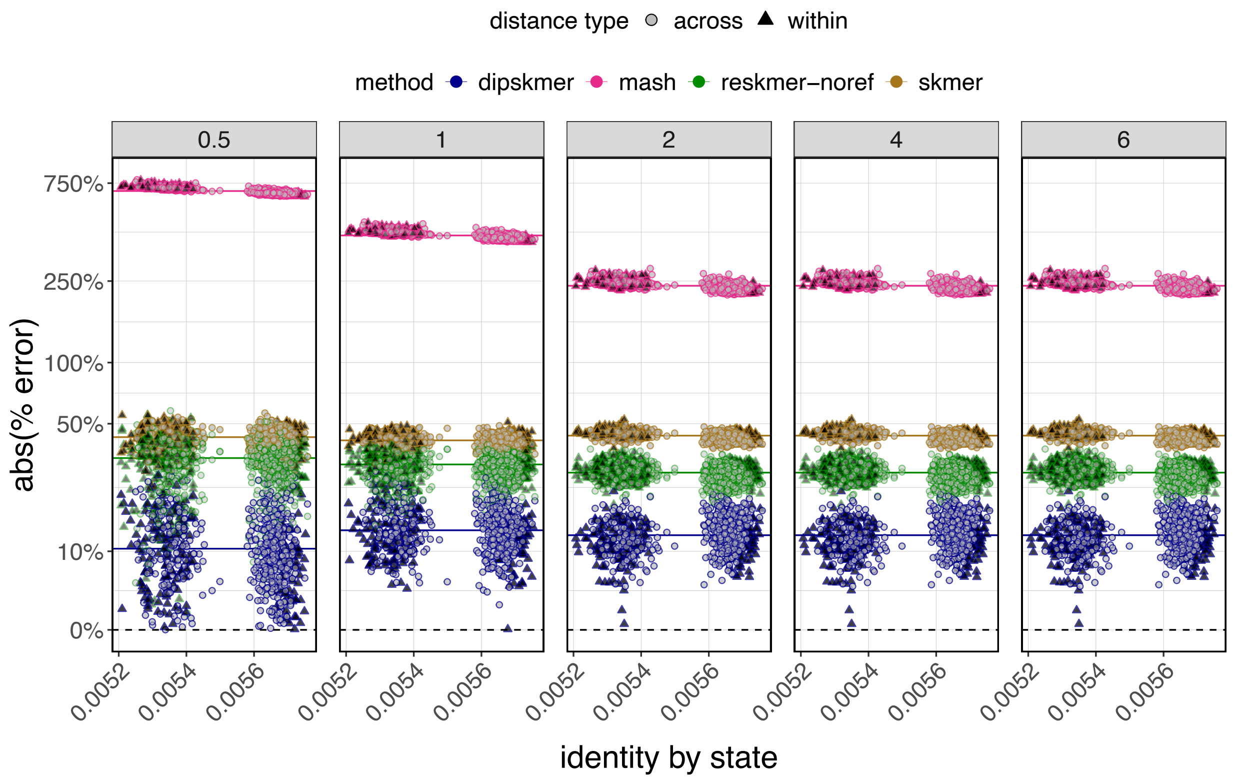


Figure S12: **Difference Between Reference-Free $\theta$ Estimates and Angsd IBS:** Results for different coverage are separated into panes (*top strip*). On the $x$-axis we plot IBS as estimated by Angsd. Some outlier datapoints with very small IBS distances are removed for ease of visualization (See Fig. 5A). Solid horizontal lines point to the mean % error for each method.

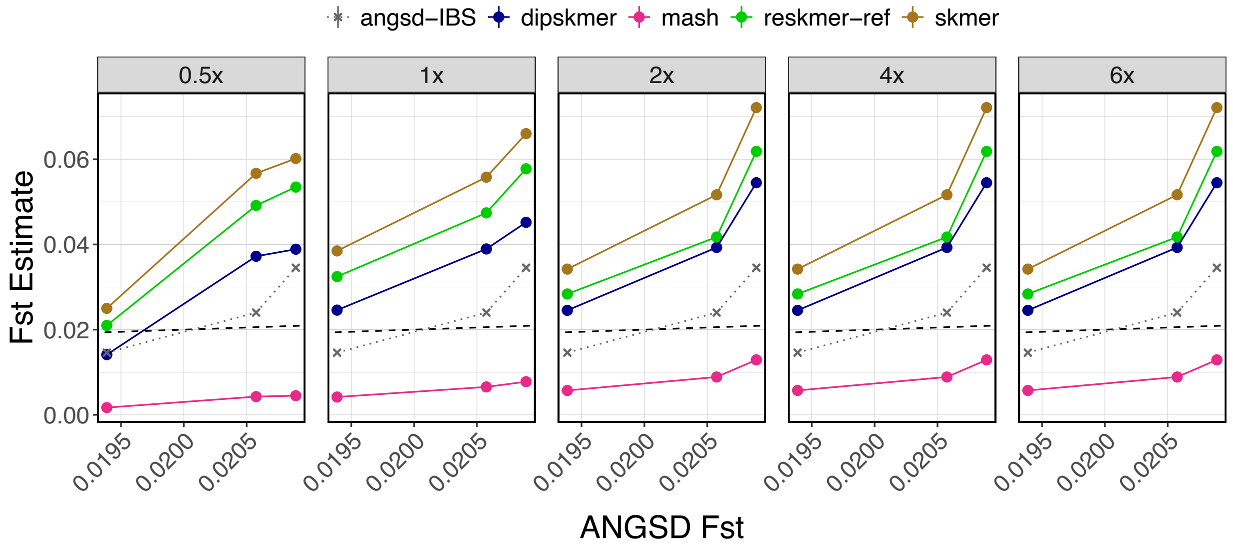


Figure S13: **Comparing Reference-Free $F_{st}$ Estimates to Angsd $F_{st}$ obtained via 2D-SFS:** Dotted line represent the $F_{st}$ used in the $x$-axis in Fig. 5C. The $x$-axis is Angsd's default $F_{st}$ estimator using 2D site frequency spectra (2D-SFS).
